# Kingdom-specific lipid unsaturation shapes up sequence evolution in membrane arm subunits of eukaryotic respiratory complexes

**DOI:** 10.1101/2024.07.01.601479

**Authors:** Pooja Gupta, Sristi Chakroborty, Arun K. Rathod, Shreya Bhat, Suparna Ghosh, Pallavi Rao T, R Nagaraj, Moutusi Manna, Swasti Raychaudhuri

## Abstract

Sequence evolution of protein complexes (PCs) is constrained by protein-protein interactions (PPIs). PPI-interfaces are predominantly conserved and hotspots for disease-related mutations. How lipid-protein interactions (LPIs) constrain sequence evolution of membrane- PCs? We explore Respiratory Complexes (RCs) as a case study as these allow to compare sequence evolution in subunits exposed to both lipid-rich inner-mitochondrial membrane (IMM) and aqueous matrix. We find that lipid-exposed surfaces of the IMM-subunits but not of the matrix subunits are populated with non-PPI disease-causing mutations signifying LPIs in stabilizing RCs. Further, IMM-subunits including their exposed surfaces show high intra-kingdom sequence conservation but remarkably diverge beyond. Molecular Dynamics simulation suggests contrasting LPIs of structurally superimposable but sequence-wise diverged IMM-exposed helices of Complex I (CI) subunit Ndufa1 from human and *Arabidopsis* depending on kingdom-specific unsaturation of cardiolipin fatty acyl chains. *in cellulo* assays consolidate inter-kingdom incompatibility of Ndufa1-helices due to the lipid-exposed amino acids. Plant-specific unsaturated fatty acids in human cells also trigger CI- instability. Taken together, we posit that altered LPIs calibrate sequence evolution at the IMM-arms of eukaryotic RCs.

## Introduction

Proteins interact, and often act as protein complexes (PCs)^1,2^. PCs executing fundamental cellular functions are conserved from prokaryotes to eukaryotes. Structures of eukaryotic PCs with prokaryotic ancestry are subtly refined in different kingdoms of life via alterations of assembly states and gain or loss of subunits^2–4^. Sequence of the PC-subunits simultaneously evolve but rarely diverge at the protein-protein interaction (PPI) surfaces that define the core composition, structure, and function of the complexes. Mutations of these conserved interactions are deleterious and associated with diseases^5–8^. In contrast, exposed PC-surfaces that do not engage into PPIs tend to be least conserved^9–12^. Intriguingly, lipid-protein interactions (LPIs) are implicated in the structural organization of membrane-PCs^13–17^ but how LPIs impact sequence evolution of membrane-PCs is insufficiently addressed.

Five respiratory complexes (Complex I to V; RCs) are fundamental for ATP production in aerobic eukaryotes. Nuclear-encoded RC-subunits translocate from cytoplasm, assemble with mitochondria-encoded subunits at the inner-mitochondrial membrane (IMM) and mitochondrial matrix, and perform oxidative phosphorylation (OxPhos)^18–21^. All the RCs contain matrix and IMM-arms with subunits exposing surfaces to both hydrophilic matrix and hydrophobic lipids. Mutations accumulate much faster in mito-genome than nuclear-genome and mutation-rates can even vary between two close species within the same eukaryotic kingdom^22–26^. Nuclear-encoded RC-subunits co-evolve to compensate the increased mutations in the mito-encoded counterparts at the IMM-arms resulting in interspecies incompatibility of nuclear and mito-encoded subunits^27–29^. RCs also adopt kingdom-specific accessory subunits to accommodate exclusive functions, heterogeneous cristae morphologies, and IMM lipid- chemistry across eukaryotes^30–33^. The dynamic PPIs and LPIs originated thereby at the matrix-arm and IMM-subunits of RCs provide an opportunity to compare PPI and LPI- mediated sequence evolution.

Here, we map human OxPhos-deficiency mutations on RC-subunits on the high-resolution structures of the complexes. We find that while point mutations in matrix-arm subunits destabilize intra or inter-subunit PPIs, ∼ 50% of the mutations in IMM-arms locate beyond PPI-interfaces. Rather, these non-PPI mutations populate the exposed surfaces of IMM-subunits, which display remarkable sequence divergence between eukaryotic kingdoms. Co-evolution of mito-encoded and nuclear-encoded subunits due to mito-genome instability fail to explain this phenomenon as the cross-kingdom sequence divergence is consistent in lipid-embedded mitochondrial membrane PCs lacking mito-encoded subunits. Our results indicate that kingdom-specific unsaturation in cardiolipin is an important determinant of lipid-affinity of trans-IMM helices of RC-subunits to mediate stable assembly of the complexes. Together, we posit that while surfaces of matrix-arm subunits of RCs may have evolved neutrally, surfaces of IMM-subunits undergo adaptive co-evolution along with the IMM-lipids across eukaryotes.

## Results

### Disease-related mutations populate membrane arm surfaces of respiratory complexes

Respiratory complex I (CI) is the first and largest complex in the respiratory chain ^34,35^. Human CI is composed of 44 subunits with 17 subunits in the matrix arm, 23 subunits embedded into inner mitochondrial membrane (IMM), and 4 subunits exposed to the inter- membrane space (IMS) (**Fig. 1A**). Fourteen of these subunits are designated as core subunits as these are conserved from bacteria to modern eukaryotes and distributed equally between the matrix and IMM-arms. Others are eukaryotic innovations and known as accessory subunits. OMIM database indicated 131 mutations in CI-subunits associated with OxPhos- deficiency diseases including Leigh, LHON, and MELAS Syndrome (**Fig. 1B, Table S1A**).

**Fig 1:**
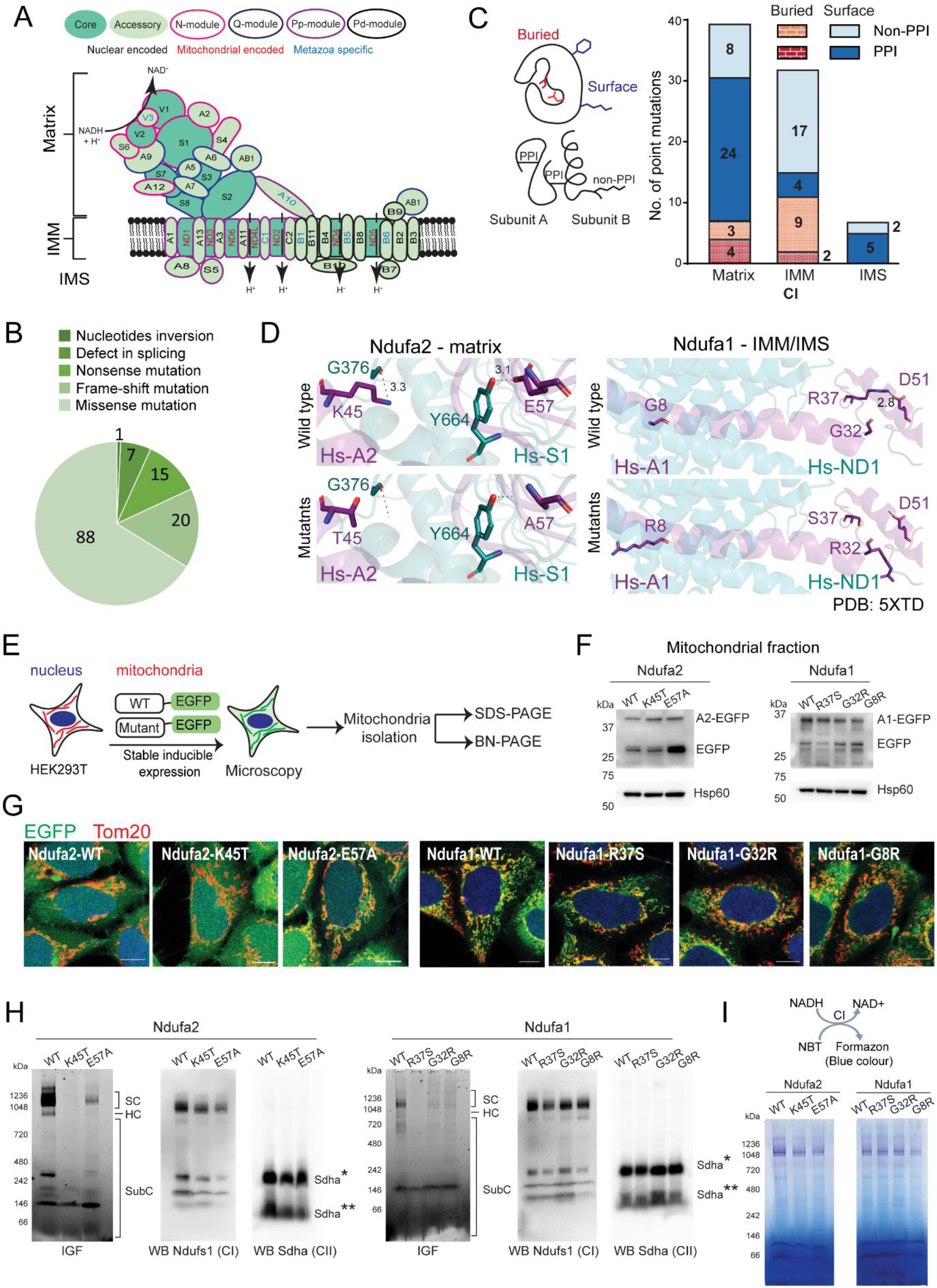
Mutations causing CI-deficiency diseases are populated at the surface of membrane arm subunits. A. Schematic representing structural framework of metazoan CI. 44 subunits associate to form 4 functional modules namely NADH dehydrogenase module (N-module; pink), ubiquinone reduction module (Q-module; blue), proton pumping proximal and distal modules (PP and PD modules; purple and black). Curved arrow: reaction catalysed by N-module; straight arrows: flow of protons (H^+^). NAD^+^ - Nicotinamide adenine dinucleotide; NADH - Nicotinamide adenine dinucleotide hydrogen. Nuclear encoded and mitochondria encoded subunits to be prefixed as Nduf and Mt, respectively. B. Pie chart indicating the different types of mutations associated with CI-deficiency diseases including MELAS, Leigh and LHON syndrome as obtained from OMIM database (omim.org). C. **Left:** Cartoon explaining buried and exposed amino acids. Involvement in protein-protein interactions (PPI) or otherwise (non-PPI) indicated. Amino acids with no neighbouring atom in 3 Å^2^ space were considered as surface exposed. PPI includes hydrogen bond, salt bridge, cation-π interaction. **Right:** Stacked bar graph indicating the distribution of the 78 missense CI-deficiency mutations as observed on CI- structure (PDB: 5XTD) into various categories as mentioned in left schematic. D. Stick representation of PPI and non-PPI amino acids in Ndufa2 and Ndufa1 as visualized by PyMol. Ndufa2 and Ndufa1: purple; interacting subunits: teal. Dashed black lines indicating PPI with bond length in Å. E. Schematic representing the workflow used for fig F-I. F. SDS PAGE followed by western blots indicating levels of wild type (WT) and mutant proteins in mitochondrial fraction probed with EGFP antibody. Hsp60 as loading control. G. Microscopy images showing EGFP tagged WT and mutant subunits. Each image represent single Z-section of 0.3 μm. DAPI (blue): nucleus. Tom20: mitochondrial marker. Scale bar: 10 μm. H. *in-gel* EGFP fluorescence (IGF) assays of overexpressed wild type and mutant CI-subunits and western blots (WBs) of digitonin solubilised mitochondrial fractions of the same probed for CI and CII subunits using anti-Ndufs1 and anti-Sdha. SC- supercomplex; HC- holocomplex; SubC- subcomplex. Sdha*- HC; Sdha**- SubC. I. *in-gel* CI activity assays. **Top:** Reaction scheme. **Bottom:** BN-PAGE gels processed for CI activity.

Out of these, 43 mutations represented nucleotide-inversions, splicing defects, nonsense mutations, and frameshifts resulting in either truncation or extensive modulation of subunit sequences. Rest 88 missense mutations change single amino acids of which 7 showed positional-discrepancy as per assignment in the highly resolved cryo-electron microscopy structure of human CI^36^, 2 were redundant reports, and 1 mutation was at the mitochondria targeting signal of the nuclear encoded subunit Ndufa10 (**Table S2A**). We mapped the rest 78 point mutations on the solved structure of CI to identify the pivotal PPIs defining the PC.

Thirty-nine mutations were mapped on the matrix arm (**Fig. 1C, Table S2A**). Seven of these involved buried amino acids (atoms found within 3Å^2^ of neighboring space). Four buried amino acids were found to be engaged in either inter-subunit or intra-subunit inter-chain H-bonding, salt bridge, or cation-π interactions (**Fig. 1C and Table S2A**). Two mutations, A159D in Ndufs8 and P76Q in Ndufb8, switches the amino acids from nonpolar to polar. V122M in Ndufs7 may result in steric hindrance with neighboring C171, L134, S149 and M120. Among the 32 matrix-arm mutations that involved amino acids with at least 1 atom exposed (no neighboring atoms within 3Å^2^ space), 24 were engaged in PPI. Seven mutations were mapped on the IMS-subunits and 5 of these participate in intra-subunit inter-chain PPI in the solved structure (**Fig. 1C and Table S2A)**. Thus, majority of the matrix arm and IMS mutations were likely to destabilize PPIs. In contrast, 9 and 17 mutations out of the total 11 and 21 mutations that involved buried and exposed amino acids at the IMM-arm were not engaged in PPI (**Fig. 1C, Table S2A**). Non-PPI mutations were also common at the exposed amino acids on the IMM-arms of other RCs (**Fig. S1A, Table S1B-E, S2B-E**).

We focused on candidate matrix-subunit Ndufa2 and IMM-subunit Ndufa1 to investigate CI-instability in presence of PPI and non-PPI mutations. As per human CI- structure (PDB:5XTD)^36^, these subunits interact with least number of neighboring subunits in the matrix and IMM-arm respectively and thus contain both PPI-interfaces and extended exposed surface stretches. Ndufa2 is a 99 amino acid protein with 2 α-helix and 4 β-strands and interacts with Ndufs1. K45 at the 2^nd^ helix and E57 just outside the 2^nd^ β-strand were found to be mutated in diseases^37^. Both the amino acids were within H-bond distance with G376 and Y664 of neighbouring subunit Ndufs1 and the disease-mutations K45T and E57A destabilized these PPIs (**Fig. 1D**). N-terminal IMM-helix containing subunit Ndufa1 interacts with ND1 within the IMM and with Ndufa8 at the IMS. Three mutations are reported in Ndufa1^38^. R37S at the IMS-exposed C-terminal of Ndufa1 was found to be destabilizing intra-subunit H-bonding with D51 (**Fig. 1D**). G32 is also IMS exposed but close to IMM- surface, do not participate in PPI, and found to expose the arginine side chain after mutation (**Fig. 1D**). The other mutating amino acid G8 is positioned at the N-terminus of the trans- IMM helix, not involved in any PPI but likewise G32 exposes a +vely charged side-chain when mutated to arginine in disease (**Fig. 1D**).

Knockout of Ndufa1 and Ndufa2 in HEK293T cells structurally and functionally destabilize CI. Knockout cells are OxPhos-inefficient, and poorly proliferating^39^. We improvised an *in cellulo* substitution strategy to assay CI-instabilities due to the mutations in Ndufa1 and Ndufa2 in steady-state OxPhos-efficient cells^40^ (**Fig. 1E**). We overexpressed EGFP-tagged Ndufa1 and Ndufa2 which entered mitochondria (**Fig. 1F-G and S1B**) and competitively replaced the endogenous subunits in CI-assemblies without affecting stability and function of the complex as evident by *in-gel* EGFP fluorescence assays, western blots for endogenous CI-subunit Ndufs1 and CII-subunit Sdha, and *in-gel* CI-activity assays (**Fig. 1H- I and S1C-D**). Disease-related mutants of Ndufa1 and Ndufa2 also localized to mitochondria (**Fig. 1F-G and S1B**). Endogenous CI-assemblies and CI-function were not perturbed in the mutant expressing cells but unlike the wild type subunits, the mutants could not efficiently substitute the endogenous counterparts in *in-gel* fluorescence assays despite no perturbations of their predicted tertiary structures after mutations (**Fig. 1H-I and S1E**). Thus, likewise the PPI-mutants of Ndufa1 (R37S) and Ndufa2 (K45T and E57A), non-PPI mutants of Ndufa1 (G8R and G32R) were also outcompeted by the endogenous subunits due to their structural incompatibilities within *in cellulo* CI-assemblies. We posit that while steric hindrance can explain destabilization of the CI-structure due to non-PPI mutations at the buried hydrophobic core of the PC, altered interactions with the IMM-lipids might cause CI- instability due to mutations at the non-PPI amino acids at the surface.

### IMM-arm subunits are sequence-conserved within eukaryotic kingdoms but diverged beyond

Next, we investigated the conservation of disease-related amino acids across eukaryotes. Both the disease-causing PPI mutations K45T and E57A on the CI matrix arm subunit Ndufa2 were conserved between metazoans, Fungi, and plants (**Fig. 2A**). Similarly, 21 out of the 27 amino acids that are involved in PPI-mutations on various subunits of CI matrix-arm were found to be conserved (**Fig. 2B, Table S2A and S2F)**. Six disease-related but unconserved matrix-arm amino acids were also involved in PPI (**Fig. 2B**). Amino acids in the same positions in *A. thaliana* and *Y. lipolytica* CI-structures (PDB:7ARB and 7O71)^41,42^ were either absent or not resolved, non-interacting, or participating in different interactions suggesting subtle micro-arrangement of PPI in the CI matrix-arm in different kingdoms of eukaryotic life (**Fig. S2A**).

**Fig 2:**
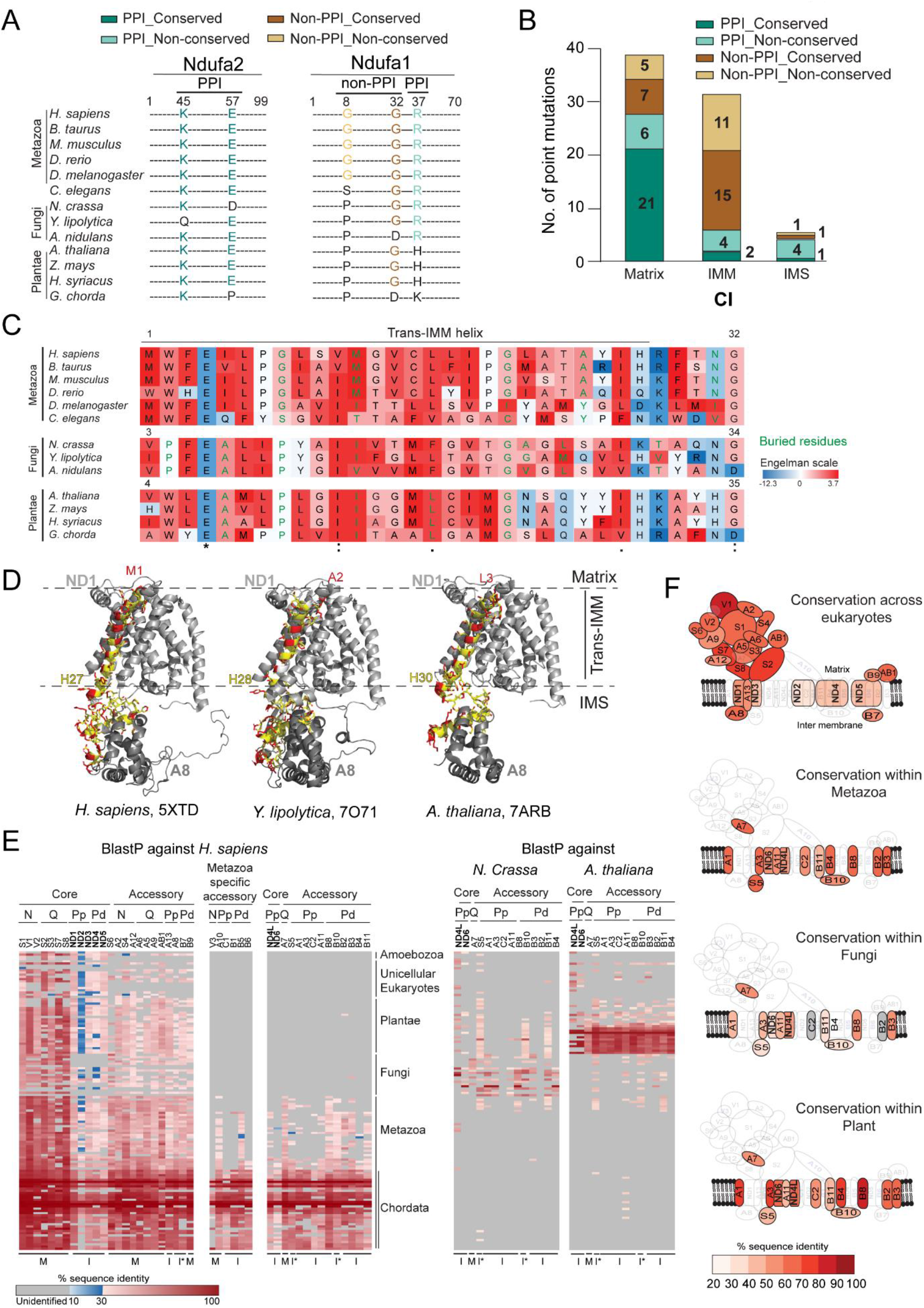
IMM-arm subunits of CI are sequence conserved within eukaryotic kingdoms A. Multiple sequence alignment indicating conservation of amino acids of Ndufa2 and Ndufa1 mutated in human CI-deficiency diseases. Sequences of 13 representative organisms from Metazoa, Fungi and plants aligned. B. Stacked bar graph representing conservation of 78 missense mutation involved in CI-deficiency diseases. Sequences of 13 representative organisms from Metazoa, Fungi and plants indicated in **2A** analyzed. C. Conservation and hydrophobicity of trans-IMM helix of Ndufa1 is represented. Color coded as per Goldman-Engelman-Steitz (GES) hydrophobicity scale indicating packing ability of trans-IMM helix into membrane. Black font: surface exposed amino acids; green font: buried amino acids. 1^st^ and last amino acid indicated with number at top of table of each kingdom. *– identical; : – strongly similar; . – weekly-similar as per Clustal Omega convention. D. Nudfa1 structure along with neighbouring subunits from CI-structures of *Homo sapiens* (PDB: 5XTD)*, Yarrowia lipolytica* (PDB: 7O71) *and Arabidopsis thaliana* (PDB: 7ARB) representing Metazoa, Fungi and plants. Red: surface exposed atoms and amino acids; yellow: partially exposed atoms and amino acids to surface; white: buried amino acids. The initial and end amino acid of IMM helix is indicated by single letter amino acid code followed by amino acid position in sequence. Light grey cartoon: neighbouring IMM subunit ND1; dark grey cartoon: neighbouring IMS- subunit Ndufa8. E. Heatmaps representing % sequence identity of CI-subunits when searched using blastp against organisms mentioned. 134 organisms from eToL were considered for analysis. Sequences of *Homo sapiens, Neurospora crassa and Arabidopsis thaliana* from Metazoa, Fungi and plants were used for blast search. Grey box indicate no blastp hit found. M - matrix; I - IMM; I* - IMS; C - Core; A - Accessory F. Schematic representation indicating the mean % identity of CI-subunits calculated from **2E**.

PPI micro-arrangement was more prominent in the IMM-arm of CI as 4 out of 6 disease-related amino acids involved in PPI in human CI were unconserved (**Fig. 2B, Table S2A and S2F)**. Three of these were on the mito-encoded core subunits suggesting evolving pivotal interactions with neighbouring nuclear-encoded subunits (**Table S2A**). Intriguingly, divergence of disease-related PPIs was also observed in other RCs including CII, which lacks mito-genome encoded subunits (**Fig. S2B, Table S2B-E and S2G-J**). This suggested that evolution of PPIs at the IMM-arms of RCs might not be fully explained by increasing mutations rates in mito-encoded subunits.

Interestingly, 15 conserved non-PPI amino acids on the IMM-subunits were also mutated in CI-destabilizing diseases (**Fig. 2B**). These mutants were found to expose charged or bulky side chains suggesting that physicochemical interactions between these conserved amino acids with IMM-microenvironment could be pivotal in stabilizing CI (**Fig. S2C and Table S2A**). Similarly, 1 IMS mutation G32R on conserved Glycine of Ndufa1 positioned at closed proximity of the IMM-surface was found to expose +vely charged side chain of arginine (**Fig. 1D and 2A**). We also found 11 unconserved non-PPI mutations on various IMM-subunits (**Fig. 2B**). One of these, G8 at the trans-IMM helix of Ndufa1 was otherwise a buried residue in human CI but +vely charged side chain of arginine was exposed after mutation (**Fig. 1D and 2A, C**). In plants and Fungi, we observed a proline residue in the same position (**Fig. 2A, C**). Apart from this proline, multiple amino acids along the trans- IMM helix of Ndufa1 also diverged between plants, Fungi and metazoans in terms of charge and degree of hydrophobicity (**Fig. 2C**). Ndufa1 is neighboured by ND1 at the IMM and by Ndufa8 at IMS^36^. These neighbouring subunits did not bury Ndufa1 entirely rather the trans- IMM helix of Ndufa1 was surface-exposed in Metazoa, plant, or Fungi CI indicating a possibility of evolving interactions with IMM-lipids across eukaryotes **(Fig. 2D**).

Sequence divergence of Ndufa1 trans-IMM helix prompted us to analyse sequence evolution of other IMM-subunits of CI across 134 organisms along the eukaryotic Tree of Life (eTOL)^43^ (**Fig. S2D, Table S3**). Sequences of matrix and IMS subunits were analysed as control. Using blastp, we identified sequence-homologues of all matrix-arm core subunits of human CI across all forms of eukaryotes (**Fig. 2E and 2F, Table S4A and S5A**). Sequence identity was highest (∼80%) within the metazoan homologues while plant and Fungi homologues showed ∼60% sequence identity (**Fig. S2E**). Metazoan homologues of 9 matrix-arm accessory, 5 IMM-arm core, and 3 P-module accessory subunits of human CI showed high sequence identity of ∼60% (**Fig. 2E and S2E**). In contrast, mean sequence identity of plants and Fungi homologues of the same human subunits was reduced to ∼40% (**Fig. 2E and S2E**). No blast-hits were obtained for mitochondrial encoded core subunits ND4L and ND6 beyond Metazoa.

Human CI contain 6 metazoan-specific accessory subunits distributed between N- and P-modules (**Fig. 1A and 2E**). Expectedly, sequence homologues were not identified for any of these beyond metazoans. Sequence homologs of 1 Q-module accessory subunit Ndufa7, 2 IMS, and 9 IMM-arm accessory subunits distributed between PP and PD-modules were also not identified among Fungi, plants, and unicellular eukaryotes by blastp search (**Fig. 2E**). We searched homologues of these subunits using sequences of Fungi *N. crassa* and plant *A. thaliana* that revealed homologs with 60-80% sequence identity within the respective kingdoms (**Fig. 2E and S2E**). Thus, among the IMM accessory subunits Ndufa13 was sequence conserved all across, 4 subunits were metazoan specific, and 9 subunits were uniquely sequence-conserved within eukaryotic kingdoms but extensively diverged beyond (**Fig. 2E-F and Table S4A and S5A**). Interestingly, among Fungi, sequences for most of the subunits of *Y. lipolytica* were uniquely conserved with very few homologs even within the kingdom (**Table S4B and S5B**) although the PDB structure (PDB: 7O71) is nearly superimposable with human CI (PDB:5XTD).

Similar to CI, IMM-subunits were sequenced-conserved within kingdoms and diverged beyond for all the RCs (**Fig. S3, Table S4C-F and S5C-F**). Human CII is constituted by 4 core nuclear-encoded subunits^44^ while plant CII contains 8 subunits^45^. Among the human subunits, Sdhc and Sdhd are IMM-embedded. Homologs of human Sdhd were found to be sequence-conserved within kingdom (**Fig. S3**). Human CIII is an obligate dimer composed of 10 subunits^46^. Six subunits are IMM-embedded out of which 3 are core subunits including the mito-encoded Mt-Cyb^36^. Rest 3 nuclear-encoded IMM-subunits were well conserved within the eukaryotic kingdoms while sequence-homologs were either poorly conserved or not found beyond kingdoms (**Fig. S3**). Three mito-encoded and 11 nuclear-encoded subunits constitute human CIV. Three nuclear-encoded accessory subunits are metazoan specific and IMM-embedded. Out of the rest 6 nuclear-encoded accessory subunits, Cox5A is opisthokont-specific and present at the matrix while others are IMM-embedded^47^. Sequence homologs of these 5 IMM-embedded accessory subunits were found within the kingdoms but poorly beyond (**Fig. S3**). Like CI, human CV is composed of well-defined matrix and IMM arms^48^. All F1 subunits and 3 FO subunits are matrix-exposed. Ten FO subunits are IMM-embedded out of which 2 are mito-encoded, 2 are metazoan specific and 2 are opisthokont-specific. Rest 4 nuclear-encoded IMM-embedded accessory subunits of human CV did not have sequence-homologs beyond metazoans while these were sequence conserved within Fungi and plants (**Fig. S3**). Remarkably, 3 matrix-exposed subunits Atp5pb, pd, and f6 were similarly sequence-conserved within the eukaryotic kingdoms but not beyond. These 3 subunits constitute the FO peripheral stalk as a connecting bridge between F1 and F ^48^.

We also analysed the subunit-sequences of mitochondrial outer membrane translocase Tom40 which switches between dimers and trimers^49^. Seven proteins constitute the complex which are either fully or partially OMM-embedded. All these proteins were sequence-conserved within the eukaryotic kingdoms. Sequence homologs were either not found by blastp search beyond the kingdoms or if found, sequence homology was below 30% (**Fig. S4, Table S4G and S5G**). Both CII and Tom40 complex are devoid of mito-encoded subunits but still showed consistent kingdom-specific sequence conservation. This supports our assumption that in addition to the diverging mito-genome, sequence evolution of membrane- subunits of mitochondrial PCs could also be contributed by membrane lipids.

### Trans-IMM helices of Human and Arabidopsis Ndufa1 interact differently with IMM-lipids

To consolidate the hypothesis of lipid-driven sequence evolution of IMM-subunits of RCs, we performed amino acid wise conservation analysis for the lipid-exposed regions of 8 single-pass IMM-arm accessory subunits of CI containing 25-residue continuous stretch with more than 80% exposed amino acids (**Table S6A**). These 8 accessory subunits were present in all 3 kingdoms of eukaryotic life and was well resolved in available cryo-EM structures^41–43^. As control, we also analysed similar matrix-exposed stretches of 8 matrix-arm subunits (**Table S6B**). **Fig 3A** shows % identity for each amino acids of the matrix-exposed stretch of Ndufa2 and the exposed trans-IMM helix of Ndufa1 with mean identity of 40.92% and 45.64% across eukaryotes, respectively. Kingdom-specific sequence identity for matrix- subunit Ndufa2 was restricted to 48-56% within Metazoa and plants. In contrast, exposed trans-IMM helix of Ndufa1 showed 63.64% sequence identity within Metazoa and 67.52% within plants. Consistently, exposed surfaces of all the 8 IMM-arm subunits showed significant increase in mean sequence identities within kingdoms (**Fig. 3B**). This suggested that the exposed surfaces of matrix-arm CI-subunits could have evolved neutrally while exposed surfaces of IMM-subunits were under kingdom-specific selection pressure possibly due to interactions with IMM-lipids.

**Fig 3:**
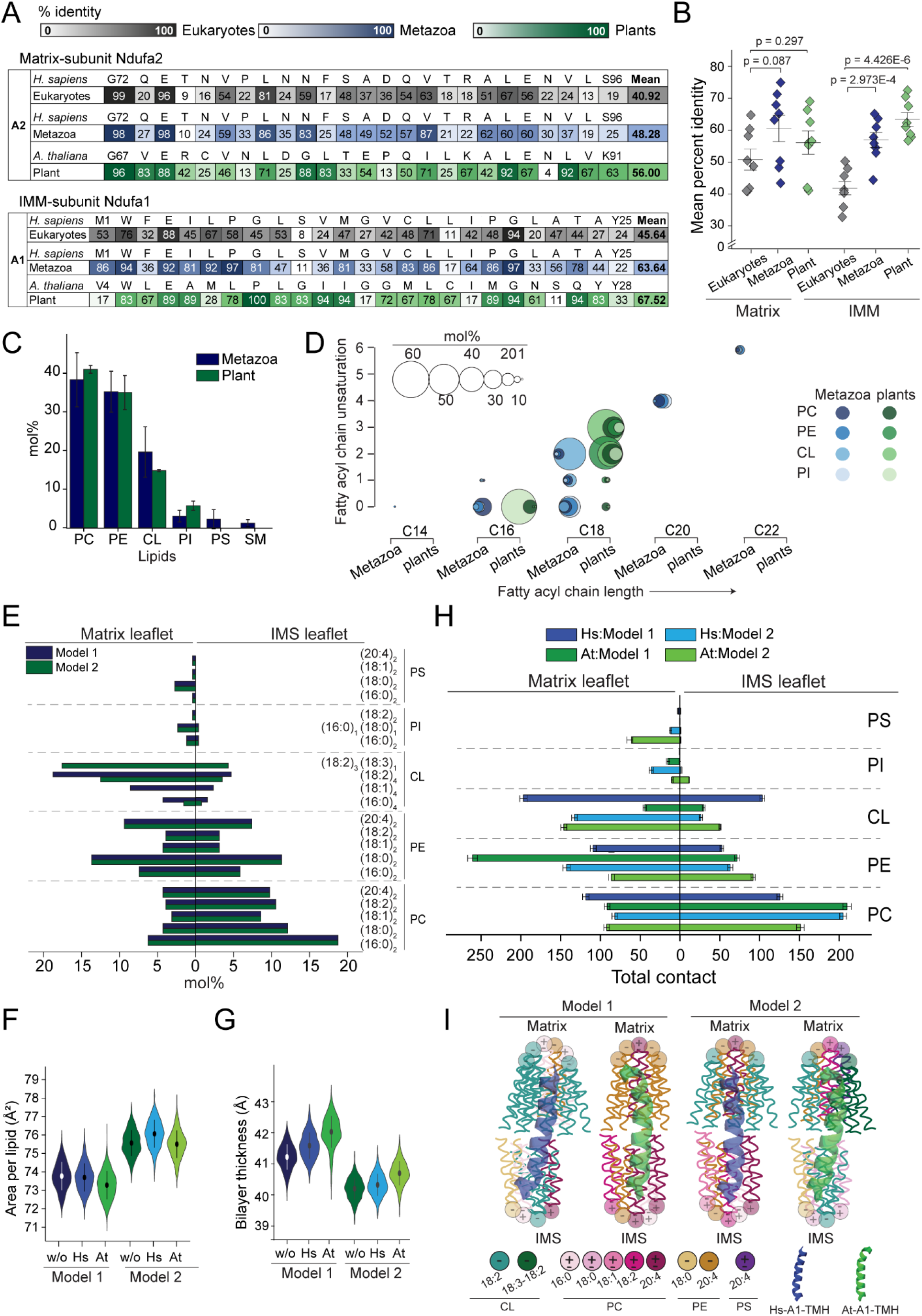
Trans-IMM helix of Human and Arabidopsis Ndufa1 interacts differently with IMM-lipids A. % sequence identity of individual amino acids at surface exposed stretches of Ndufa2 and Ndufa1. *Homo sapiens* sequences were used as reference sequence to compare multiple sequence alignment across eukaryotes and within Metazoa. *Arabidopsis thaliana* sequences were used for plants. Amino acids are represented by single letter code. 1^st^ and 25^th^ amino acids represented with number indicating the positions of amino acid in *H. sapiens* and *A. thaliana* sequences. B. % sequence identity of individual amino acids at surface exposed stretches of 8 matrix and IMM arm subunits plotted. Whisker: 1 Standard deviation from the mean. Two sample student t-test. Details in **Table S6A-B**. C. Bar graph comparing mol% of different lipids in IMM between metazoa and plants. Literature reported mol% values were averaged and plotted. Error bars indicate SD. Details in **Table S7A**. D. Bubble graph showing mol% of different fatty acyl chains found in IMM. Literature reported mol% values were averaged and plotted. Details in **Table S7B**. Graph was prepared using Visual Paradigm (https://online.visual-paradigm.com/). E. Graph showing mol% of various lipids used for simulating model 1 and model 2. Literature reported mol% values were averaged and plotted. Details in **Table S7A-C, S8A-B**. F. Violin plot indicating area per lipid (Å^2^) calculated using final 500 ns of simulation time. w/o - without trans-IMM helix; At – At-A1 trans-IMM helix; Hs - Hs-A1 trans- IMM helix. Hollow circles: Mean; Error bars: 1 SD from mean. G. Violin plot indicating bilayer thickness (Å) calculated using final 500 ns of simulation time. w/o - without trans-IMM helix; At – At-A1 trans-IMM helix; Hs - Hs-A1 trans- IMM helix. Hollow circles: Mean; Error bars: 1 SD from mean. H. Graph showing time-averaged total acyl chain contacts of different lipid species of matrix and IMS leaflets with Ndufa1 trans-IMM helices i.e., human M1-M12, *A. thaliana* V4-I15 and human G13-T23, *A. thaliana* G16-Q26, respectively. Total contacts averaged over final 500 ns of simulation time. Error bar: SD. I. Schematic model summarizing interactions of acyl chains of different lipids with Human (Hs) and *A. thaliana* (At) Ndufa1 trans-IMM helix. Data obtained from **3H and S5E**. 1 lipid with chains represents 50-110 contacts; 2 lipids – 110-150 contacts; 3 lipids – 150-200 contacts. TMH – Transmembrane helix

Cardiolipin (CL) is a unique lipid present only in mitochondria membranes. CL has been implicated in the biogenesis of OMM import channel Tom40^50^. Cryo-EM structures of CI, CIII, and CIV from various species have identified IMM-lipids including CL associated with the complexes^36,41,42,47,51^. Simulation of CIII and CIV in model membranes identified CL to be the major interacting lipid^52–54^. Interestingly, mol% composition of IMM-lipids was found to be similar between plants and Metazoa (**Fig. 3C, Table S7A**) but the fatty acyl chain length and unsaturation were uniquely kingdom-specific (**Fig. 3D, Table S7B**). CL is reported to be more unsaturated in plants (18:3 and 18:2, 55:38) than Metazoa (18:2 and 18:1, 49:20) while 20:4 and 22:6 phosphatidylcholine (PC), phosphatidylethanolamine (PE), and phosphatidylinositol (PI) are reported in Metazoa only^32,55^ (**Fig. 3C-D, Table S7B**). Also, sphingolipids, cholesterol, and phosphatidylserine (PS) are not reported in plant mitochondria membranes till date (**Fig. 3C and Table S7A**).

We simulated a model IMM mimicking Metazoa lipid composition (Model 1, **Fig. 3E**). Variation of lipids between IMM-leaflets in Metazoa^56–58^ was considered with PI, PE, and CL rich on matrix and PC rich on IMS leaflets (**Fig. 3E, Table S7C**). Low abundant IMM-lipids sphingomyelin, sterols, lysophosphatidylcholine, lysophosphatidylehtanolamine, phospahtidylglycerol etc. were not included for simulation. Fatty acids 16:0, 18:0, 18:1, 18:2, and 20:4 were simulated as per their abundance in metazoan IMM. We also simulated a Model 2 where we introduced plant-specific unsaturation 18:3 at sn1 position of 1^st^ phosphate of otherwise 18:2 CL, which resulted in 18:3-18:2, 18:2, and 16:0 CL in 55:40:5 ratio. Other lipids in Model 2 remained identical to Model 1 (**Fig. 3E, Table S8A**). Total 256 lipid molecules were placed in each leaflet and all-atom MD simulations were performed for 1 μs **(Table S8B)**.

Replacing just one acyl chain from 18:2 to 18:3 in CL in Model 2 resulted in alteration in membrane structural properties. Area per lipid (APL) in Model 2 was increased to 75.6 Å^2^ from 73.7 Å^2^ in Model 1 (**Fig. 3F and S5A**). The mean bilayer thickness of Model 1 was 41.2 Å while that of Model 2 was 40.2 Å (**Fig. 3G and S5B**). We then simulated trans- IMM helices of human (Hs) and *Arabidopsis* (At) Ndufa1 (Hs-A1 and At-A1) within the IMM-models. IMM-exposed helix of Hs-A1 was constituted by N-terminal M1-H27 while corresponding At-A1 helix was from V4-H30 (**Fig. 2C**). Y25-H27 in Hs-A1 was not included for simulation as the stretch was found to be identical as Y28-H30 in At-A1 (**Fig. 2C**). Y27 in At-A1 helix corresponded to A24 in Hs-A1. Y27 was found to be involved in cation-π interaction with M98 of ND1 in At-A1 but A24 was non-interactive as per the human CI PDB (**Fig S5C**). We excluded At-A1(Y27) and Hs-A1(A24) for simulation. Thus, 1^st^ 23 amino acids of Hs-A1 (M1-T23) and the corresponding (V4-Q26) in At-A1 were not involved in PPIs, were sequence diverged between kingdoms, and therefore considered for simulation. We placed single trans-IMM helices of each in the model membranes. Both the trans-IMM helices lied within the membrane normal in both model membranes indicating that they were accommodated within the membrane hydrophobic core rather than interacting with the head groups of the lipids (**Fig. S5D**). The mean APL remained unaltered when Hs- A1 trans-IMM helix (M1-T23) was placed in Model 1 while that slightly decreased to 73.2 Å^2^ with At-A1 (V4-Q26) (**Fig. 3F and S5A**). The mean bilayer thickness of Model 1 was increased to 41.5 Å for the Hs-A1 trans-IMM helix and 42.3 Å for At-A1 (**Fig. 3G and S5B**). Conversely, mean APL remained unaltered when At-A1 trans-IMM helix was placed within Model 2 while Hs-A1 trans-IMM helix marginally increased the APL to 76 Å^2^ (**Fig. 3F and S5A**). The mean bilayer thickness of Model 2 remained unaltered when Hs-A1 trans-IMM helix was placed while At-A1 trans-IMM helix increased the mean bilayer thickness to 40.7 Å (**Fig. 3G and S5B**).

Number of contacts between the peptides and the acyl chains in the model membranes are calculated considering contacts between the non-hydrogen atoms of these groups within a cut-off distance of 5 Å^59,60^. We considered N-terminal 1-12 amino acids of the trans-IMM helices as matrix leaflet interacting while the rest was considered to interact with IMS-leaflet lipids. **Fig 3H** and **S5E** indicates total contacts of the acyl chains with trans-IMM helices averaged over the final 500 ns of simulation time. At-A1(V4-I15) in Model 1 established maximum interactions with PE (260.7 contacts) of which 160.5 and 63.7 interactions with 20:4 and 18:0 PE at the matrix leaflet, respectively. Matrix leaflet interactions with CL was found to be 44.3 amongst which 41.5 interactions were with unsaturated CL18:2. On the other hand, Hs-A1 (M1-M12) established maximum contacts with CL (196.8 contacts) amongst which 181.2 was with unsaturated 18:2. Further, interaction of Hs-A1(M1-M12) with 20:4 PE at the matrix leaflet was only 61.4 in contrast to 160.5 of At-A1(V4-I15) suggesting that both the trans-IMM helices mostly interacted with unsaturated acyl chains at the matrix leaflet of Model 1 but the interacting lipids differed (**Fig. 3H-I and S5E, Table S8C**).

In Model 2, At-A1(V4-I15) established 146 interactions with CL at the matrix leaflet with 75.6 and 65.3 contacts with 18:3-18:2 and 18:2 CL. 20:4 PE interactions were reduced to 42.8 contacts compared to Model 1. Interestingly, Hs-A1(M1-M12) gained interaction with PE (142.6) in Model 2 as compared to Model 1 (109.1) and CL interactions in Model 2 was reduced to 132.5 as compared to 196.8 in Model 1. However, even in Model 2, Hs-A1(M1- M12) preferred 18:2 CL (86.4) as compared to 18:3-18:2 CL (45.3). Thus, At-A1(V4-I15) interacted more with unsaturated PE in Model 1 in absence of 18:3-18:2 CL at the matrix leaflet whereas the Hs-A1(M1-M12) interacted more with PE as compared to CL in presence of 18:3-18:2 CL in Model 2 (**Fig. 3H and S5E, Table S8C**). At the matrix leaflet both the trans-IMM helices interacted with PC as well, however, Hs-A1(M1-M12) interacted with PC more in Model 1 (118 contacts) as compared to Model 2 (81.9 contacts). At-A1(V4-I15) established similar number of contacts with PC in both the models (**Fig. 3H and S5E, Table S8C**).

Both the trans-IMM helices interacted with PC, CL and PE at the IMS leaflet. At- A1(G16-Q26) interacted with unsaturated PC (18:2 and 20:4) and PE 20:4 (51.7, 99.1 and 57.7 contacts, respectively) at the IMS leaflet of Model 1. In contrast, the same trans-IMM helix interacted with saturated 16:0 PC, 18:0 PC, 18:0 PE and unsaturated 18:2 CL (41.5,64.6, 58.8 and 32.5 contacts, respectively) in Model 2. On the other hand, Hs-A1(G13-T23) interacted with 16:0 PC, 18:0 PC, 20:4 PC, 18:1 CL and 18:2 CL (47.2, 33.8, 34.3, 38.3, 44.5 contacts, respectively) at the IMS leaflet of Model 1 in contrast to Model 2 where it gained interactions with unsaturated PC 18:1 (38.8), 18:2 (57.1), and 20:4 (58.4), respectively (**Fig. 3H and S5E, Table S8C**). Interaction of both the trans-IMM helices with PI and PS were minimal in both the models with exception where At-A1(G16-Q26) made 61.5 contacts with PS in Model 2.

In summary, structurally superimposable trans-IMM helices of Ndufa1 from two different kingdoms of eukaryotic life showed contrasting but complementary interactions with fatty acyl chains of IMM-lipids depending on the 18:2 or 18:3 chemistry of CL (**Fig. 3I**).

### Lipid-exposed amino acids prevents exchange of Hs-A1 trans-IMM helix with At-A1 trans-IMM helix in cell culture

MD simulations depicted that At-A1 helix preferred 18:3 CL over 18:2. Mammalian cells lack 18:3 CL. Therefore, we hypothesized that At-A1 helix might not stably interact with IMM-lipids in HEK293T cells and thereby might be incompatible to integrate into CI. To validate, we performed *in cellulo* assays using EGFP-tagged constructs of Ndufa1 containing either Hs or At helix. We replaced the non-PPI, trans-IMM helix part of Hs-A1 (M1-T23) with 4^th^-26 amino acids of At-A1 where V4 was mutated to methionine (Chimera- A1; Ch-A1). As per AlphaFold, the structures are superimposable and all the PPIs between Hs-A1 and its interacting partners ND1 and Ndufa8 remained unaltered (**Fig. 4A-B and S6A)**. A gain of interaction between N21 of Ch-A1 and T256 of human ND1 was also observed (**Fig. S6A)**. We prepared stable inducible lines for Hs-A1, At-A1 and Ch-A1. All these proteins colocalized with Tom20 stained mitochondria (**Fig. 4C and S6B**). Intriguingly, western blots of mitochondrial fractions revealed excess load of At-A1 and Ch-A1 as highly abundant cleaved products ∼3-kDa lighter in mass than the full-length proteins (**Fig. 4D**). We also prepared mitoplasts devoid of the mitochondrial outer membrane protein Tom40 but containing IMM and matrix proteins Ndufa13 and Hsp60, respectively (**Fig. S6C**). A prominent band for Hs-A1G8R mutant was observed in the mitoplast fraction (**Fig. 4E**) indicating successful import of the mutant into IMM while it was unable to integrate into CI (**Fig. 1H**). Interestingly, full-length At-A1 band was almost absent and Ch-A1 band was comparatively less intense in the mitoplast fractions than Hs-A1 and Hs-A1G8R (**Fig. 4E**). The cleaved products of At-A1 and Ch-A1 were still observed in the mitoplasts indicating that the breakdown of the full-length proteins could be an IMM-associated event. We created a construct deleting the trans-IMM helix in human Ndufa1 (Δ27Hs-A1). The cleaved product of Ch-A1 run in the same position with Δ27Hs-A1 in SDS-PAGE suggesting that cleavage might have happened after the trans-IMM helix (**Fig. 4F**). Local chemistry driven cleavage of peptide bonds after N and Q have been reported for multiple proteins^61–65^. At-A1 trans-IMM helix present in Ch-A1 contains both N and Q at the C-terminus (**Fig. 2C**). The cleaved band of Ch-A1 disappeared after introducing L in place of N21 (**Fig. 4G**). Interestingly, replacing Q23 with Hs-A1 analogous T23 did not prevent the cleavage of Ch-A1 (**Fig. 4G**). These results indicated that N containing trans-IMM helices of At-A1 and Ch-A1 interact inappropriately with human IMM and therefore unstable.

**Fig 4:**
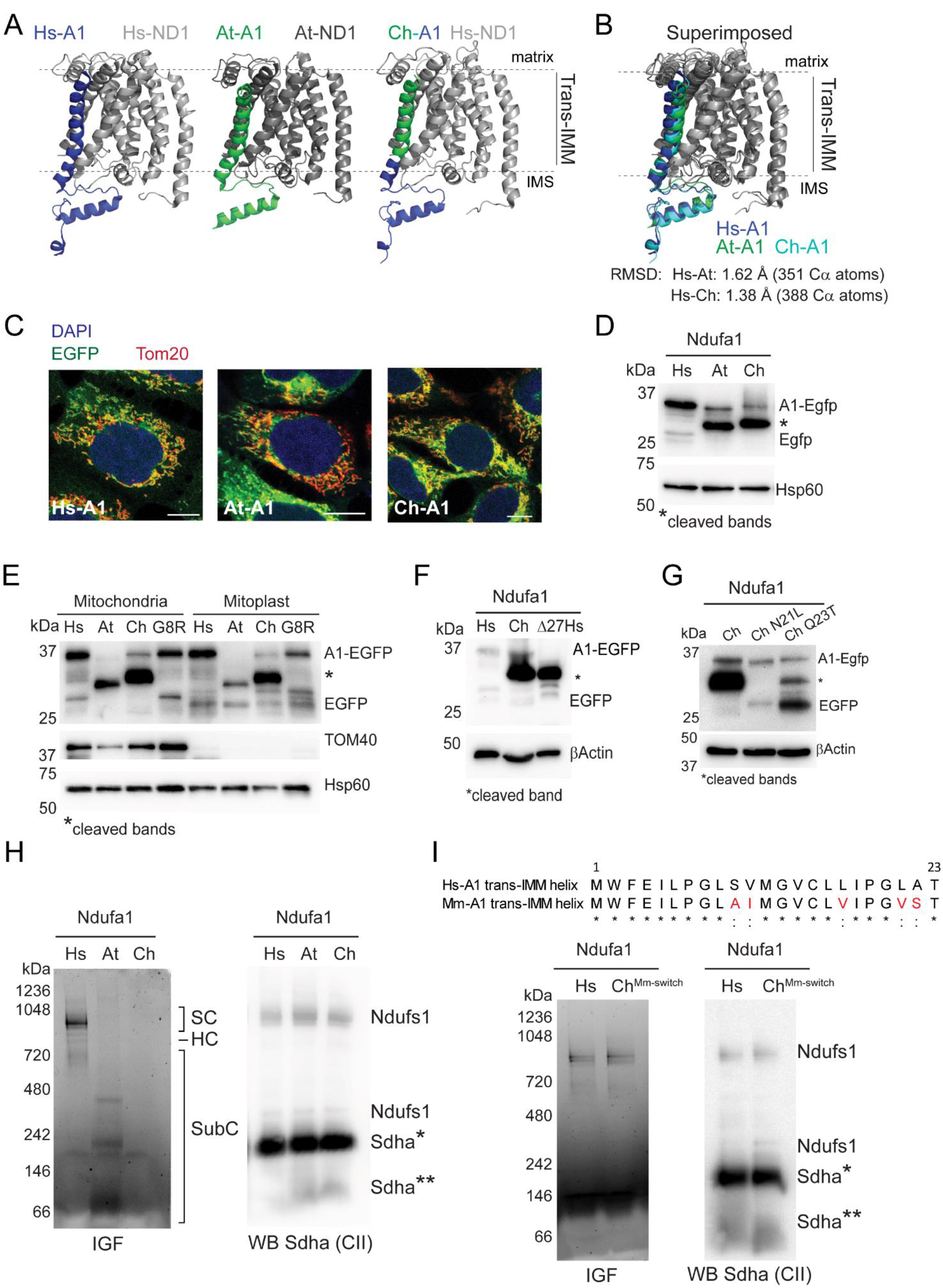
*in-cellulo* incompatibility of trans-IMM helix of At-A1 in Hs-CI A. Cartoon representations showing Ndufa1 and ND1 as visualized by PyMol. Black dotted line presents trans-IMM regions of both the proteins. **Left:** Human CI (PDB: 5XTD). **Middle:** *A. thaliana* CI (PDB: 7ARB). **Right:** Human ND1 and Chimera- Ndufa1 (Ch-A1) where trans-IMM helix of Human Ndufa1 (Hs-A1) switched with *A. thaliana* Ndufa1 (At-A1) trans-IMM helix (AlphaFold modelling). B. Superimposed images of Hs-A1 with ND1, At-A1 with ND1, and Ch-A1 with Human ND1. Black dotted line: trans-IMM regions. Hs-A1:blue; At-A1:green; Ch-A1:teal; Hs-ND1:light grey; At-ND1:dark grey. RMSD indicated are out of 388 aligned amino acids. Template alignment score: more than 0.9. C. Microscopy images of cells expressing EGFP tagged Hs-A1, At-A1, and Ch-A1. Each image represent single Z-section of 0.3 μm. DAPI (blue): nucleus. Tom20: mitochondrial marker. Scale bar: 10 μm. D. Western blot indicating Ndufa1-EGFP in mitochondrial fractions of Hs-A1, At-A1, and Ch-A1-EGFP expressing cells, probed with EGFP antibody. Hsp60 as loading control. * - Cleaved bands of A1-Egfp. E. Immunoblot indicating of Ndufa1-EGFP in mitochondrial fractions and mitoplast fractions of Hs-A1, At-A1, Ch-A1 and A1(G8R) expressing cells probed with EGFP antibody. Hsp60 as loading control. TOM40 - Outer mitochondrial membrane marker. * - Cleaved bands of A1-Egfp. F. Western blot indicating Ndufa1-EGFP in total cell lysates of Hs-A1, Ch-A1-EGFP, and Δ27Hs-A1-EGFP expressing cells, probed with EGFP antibody. Hsp60 as loading control. * - Cleaved band of Ch-A1-Egfp and full length Δ27Hs-A1-EGFP. G. Immunoblot indicating cleaved product of Ndufa1-EGFP in total cell lysates of Ch- A1, Ch-A1-N21L, and Ch-A1-Q23T expressing cells when probed with EGFP antibody. Hsp60 as loading control. * - Cleaved band of A1-Egfp. H. *in-gel* EGFP fluorescence (IGF) assay and western blot (WB) of digitonin solubilised mitochondrial fractions run on native PAGE from cells shown in **4D** and probed for CII subunit using anti-Sdha. SC - supercomplex; HC - holocomplex; SubC - subcomplex. Sdha*- HC; Sdha**- SubC. I. **Top:** Pair-wise sequence alignment of trans-IMM helix of Hs-A1 and Mm-A1. *– identical; : – strongly similar; as per Clustal Omega convention. Position of 1^st^ and 23^rd^ amino acid represented with number at top of alignment. Red font: mismatched amino acid in Mm-A1. **Bottom:** *in-gel* EGFP fluorescence (IGF) assay and western blot (WB) of digitonin solubilised mitochondrial fractions run on native PAGE from Hs-A1-Egfp and Ch - A1^Mm-switch^-EGFP, probed with anti-Sdha (CII subunit). SC - supercomplex; HC - holocomplex; SubC - subcomplex. Sdha*- HC; Sdha**- SubC.

Next, we tested if At-A1 trans-IMM helix was compatible to human CI. *in-gel* EGFP fluorescence assays using digitonin-solubilized mitochondrial extracts from Hs-A1, At-A1, and Ch-A1 cells indicated successful incorporation of Hs-A1 into CI-assemblies while both At-A1 and Ch-A1 did not integrate (**Fig. 4H**). However, two bands at ∼320 and ∼ 180 kDa was observed in At-A1 lane indicating the formation of aberrant subassemblies (**Fig. 4H**).

Endogenous RC-assemblies were not destabilized in these cells as evident by the blots with Ndufs1 and SDHA antibodies (**Fig. 4H and S6D**). CI activity was also not perturbed as per *in-gel* CI activity (**Fig. S6D**). Interestingly, we also found mismatch of five amino acids between the trans-IMM helices of human and mouse Ndufa1 (**Fig. 4I**). Therefore, we switched the first 23 amino acids of Hs-A1 with the mouse sequence (Ch-A1^Mm-switch^). The chimeric product successfully integrated into human CI indicating that intra-kingdom amino acid mismatches did not perturb LPIs of Ndufa1 (**Fig. 4I, S6E**). This is probably due to the consistency of physicochemical properties of the mismatched amino acids.

Detailed analysis of PDB:5XTD indicated that side chains of 6 out of 23 amino acids constituting Hs-A1 trans-IMM helix contained no atoms within 3 Å^2^ of neighbouring space i.e., side chains of these amino acids were entirely exposed to IMM-lipids (**Fig. 2D and 5A**). Among these, matrix leaflet F3 and P7 were conserved between human and mouse helix (**Fig. 4I**). As per simulation, F3 of Hs-A1 interacted with both PC and CL tails while P7 was more interactive with CL fatty acyl chains in model IMM 1. Interestingly, corresponding L6 and L10 of At-A1 interacted with PE chains (**Fig. 5B**). IMS-leaflet nonpolar L21 was physicochemically conserved as V21 in mouse but found to be polar as N24 in At-A1 (**Fig. 4I**). However, A22 in Hs-A1 remained physicochemically conserved as S22 and S25 in mouse and Arabidopsis helices, respectively. Simulation results indicated that Hs-A1 L21 and A22 interacted with PC and CL while corresponding N24 and S25 of At-A1 interacted with mostly PC and PE chains (**Fig. 5B**).

**Fig 5:**
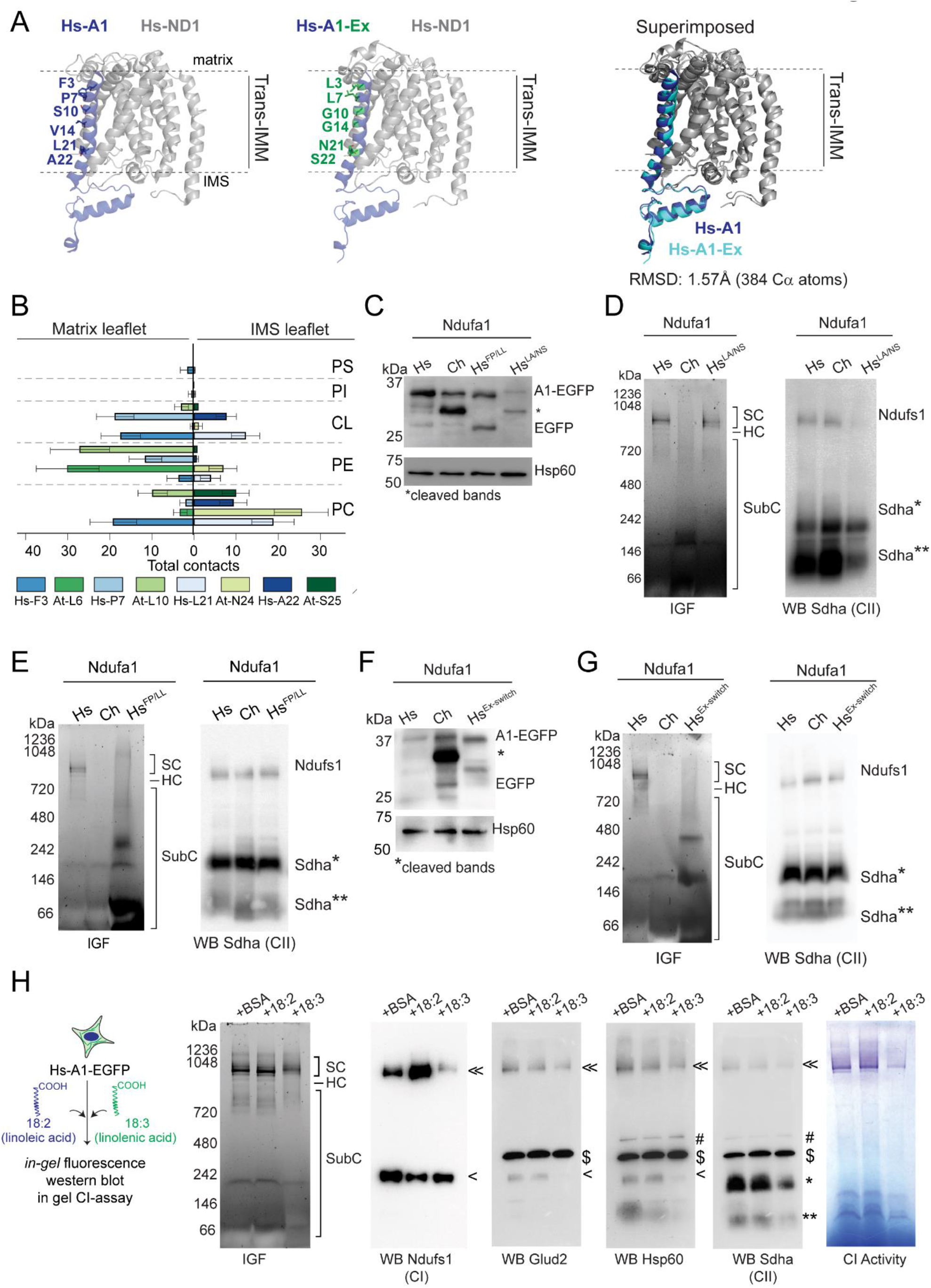
Switching lipid-exposed amino acids of Hs-A1 trans-IMM helix with At-A1 trans- IMM helix renders *in-cellulo* incompatibility A. Cartoon representations showing Ndufa1 and ND1 when visualized by PyMol. Black dotted line represents trans-IMM regions. **Left:** Ndufa1 (blue) and ND1 from human CI (PDB: 5XTD). **Middle:** Human ND1 and Hs-A1-Ex (blue-green) where exposed amino acids of Hs-A1 replaced with corresponding amino acids of At-A1 (AlphaFold modelling). **Right:** Superimposed image. Hs-A1:blue; Hs-A1-Ex:teal; Hs-ND1:light grey. RMSD indicated are out of 388 aligned amino acids. Template alignment score: more than 0.9. B. Bar graph showing total contacts made by individual amino acids of Hs-A1 and At- A1 trans-IMM helices with the acyl chains of different lipid species in model IMM1 in simulation experiment. Total contacts averaged over final 500 ns of simulation time. Error bar: SD C. Immunoblot indicating Ndufa1-EGFP in mitochondrial fractions of Hs-A1, Ch-A1, Hs-A1^FP/LL^ Hs-A1^LA/NS^ when probed with EGFP antibody. Hsp60 as loading control. *- Cleaved band of A1-Egfp. D. Native PAGE processed for *in-gel* EGFP fluorescence (IGF) assay and western blot (WB) of CII subunit Sdha using digitonin solubilised mitochondrial fractions from Hs-A1, Ch-A1 and Hs-A1^LA/NS^. SC- supercomplex; HC- holocomplex; SubC- subcomplex. Sdha*- HC; Sdha**- SubC. E. Native PAGE processed for *in-gel* EGFP fluorescence (IGF) assay and western blot (WB) of CII subunit Sdha using digitonin solubilised mitochondrial fractions from Hs-A1, Ch-A1 and Hs-A1^FP/LL^. SC- supercomplex; HC- holocomplex; SubC- subcomplex. Sdha*- HC; Sdha**- SubC. F. Immunoblot indicating Ndufa1-EGFP in mitochondrial fractions of Hs-A1, Ch-A1 and Hs-A1^Ex-switch^ expressing cells when probed with EGFP antibody. *- Cleaved band of A1-Egfp. G. Native PAGE processed for *in-gel* EGFP fluorescence (IGF) assay and western blot (WB) of CII subunit Sdha using digitonin solubilised mitochondrial fractions from Hs-A1, Ch-A1 and Hs-A1^Ex-switch^. SC- supercomplex; HC- holocomplex; SubC- subcomplex. Sdha*- HC; Sdha**- SubC. H. **Left:** Schematic representing BSA-conjugated lipid supplementation assay. **Right:** Native PAGE processed for *in-gel* fluorescence (IGF) assay, western blot (WB) using CI (Ndufs1), CII (Sdha) and other mitochondria-specific (Glud2, Hsp60) antibodies and *in-gel* CI activity. SC- supercomplex; HC- holocomplex; SubC- subcomplex. >> Ndufs1 SC; >Ndufs1 SubC; $Glud2; #Hsp60; *-Sdha HC; **- Sdha SubC.

Interestingly, when we replaced L21 and A22 with corresponding N and S from At- A1 trans-IMM helix, the resulting protein Hs-A1^LA/NS^ was cleaved and imported less into mitochondria as evident in the western blot of the mitochondrial fraction (**Fig. 5C**). However, it was efficiently integrated into human CI as evident by *in-gel* EGFP fluorescence (**Fig. 5D and S6F**). However, we did not observe any cleavage of Hs-A1^FP/LL^ where F3 and P7 were replaced with corresponding Leucine residues from At-A1 but the protein was not observed in productive CI-assemblies (**Fig. 5C and 5E)**. Instead, we observed a smear as well as a prominent but aberrant subcomplex at ∼300 kDa and faint subcomplexes at ∼600 kDa and 1048 kDa (**Fig. 5C, 5E and S6G**). Similarly, switching all the side chain exposed amino acids in Hs-A1 with the analogous At-A1 amino acids (Hs-A1^Ex-switch^) resulted in absolute disappearance of SC and HC bands in the *in-gel* fluorescence assay while two bands at ∼320 and ∼ 180 kDa indicated formation of aberrant subassemblies (**Fig. 5F-G and S6H**).

Finally, we tested if human CI could withstand engineered changes in unsaturation of fatty acyl chains in IMM-lipids. Recently, multiple works from Oemer et al. demonstrated modulation of molar composition and unsaturation of various lipids including CL in mammalian cells treated with different fatty acids during culture^66,67^. Increased incorporation of 18:2 fatty acyl chains in CL has also been linked to efficient OxPhos^67^. We similarly treated Hs-A1 cells with 18:2 linoleic acid and 18:3 linolenic acid. *in-gel* EGFP fluorescence assays revealed efficient entry of Hs-A1 in CI-HC and SCs in control vehicle-treated and linoleic acid treated cells but only partial entry was observed in linolenic acid treated cells (**Fig. 5H**). Endogenous assemblies of CI and CII were also found to be perturbed in linolenic acid treated cells as probed by Ndufs1 and SDHA western blots (**Fig. 5H**). This indicated instability of human RCs in presence of plant specific 18:3 CL. Together, these results suggested that the chemistry of the exposed amino acids in the trans-IMM helix of Ndufa1 with the surrounding saturated or unsaturated lipids determined the integration of the subunit in CI assemblies in different forms of eukaryotic life (**Fig. 6**).

**Fig 6:**
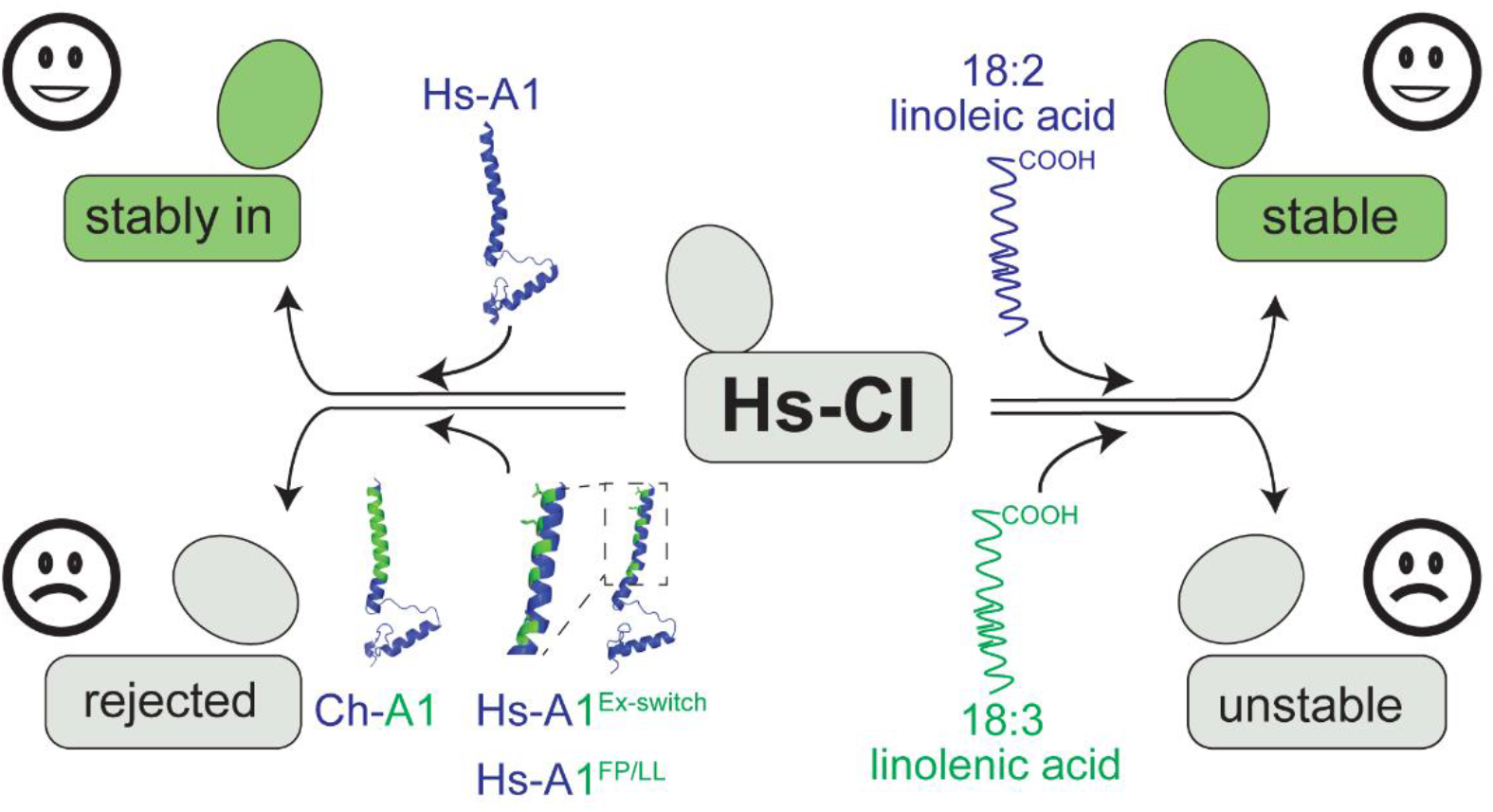
Schematic summary of *in cellulo* substitution assays. **Top left:** Human Ndufa1-EGFP (Hs-A1; blue) stably integrate into CI in HEK293T cells. **Bottom left:** Chimera Ndufa1-EGFP (Ch-A1) – Hs-A1 trans-IMM helix exchanged with At- A1, Hs-A1^FP/LL^ – Hs-A1 with lipid exposed amino acids F3 and P7 exchanged with At-A1 amino acid L3 and L7, Hs-A1^Ex-switch^ – Hs-A1 with lipid exposed amino acids exchanged with At-A1 could not stably integrate into CI holocomplex and supercomplexes. **Top right:** treating cells with 18:2 fatty acid (Linoleic acid) do not perturb CI. **Bottom right:** treating cells with 18:3 fatty acid (Linolenic acid) destabilizes CI.

## Discussion

Intra- and inter-protein interactions are established determinants of protein sequence evolution. Local chemistry with other interacting macromolecules also impose sequence divergence in proteins. Rapid evolution of mt-rRNA due to mito-genome instability is correlated with sequence-divergence of mito-ribosome^68^ and respiratory complex (RC) subunits^25^. Alteration of lipid chemistry also calibrate lipid-protein interactions^13–17^ but lipid-influenced sequence divergence of proteins is not investigated extensively. Rather, it is proposed that the sequences of membrane proteins, especially the transmembrane regions, are highly constrained by membrane lipids and therefore diverge less than other cellular proteins^69,70^. Our results indicate that membrane-proteins of mitochondrial complexes are highly sequence conserved within eukaryotic kingdoms but undergo divergent evolution across eukaryotes correlating with the evolving chemistry of surrounding lipids. In case of RCs, we posit that pivotal PPIs are evolutionarily conserved for the structural maintenance of both membrane-embedded and extra-membrane arms of these complexes. Conversely, divergence of lipid-protein chemistry at the IMM-arm is required to preserve the nearly super-imposable structural designs for optimized OxPhos across eukaryotes. We further hypothesize that higher-mutation rates in mito-genome may provide an indirect sampling advantage for protein sequences of IMM-subunits in adapting the diverging lipid environments. These hypotheses are in line with previous postulates which suggest that lipid biosynthesis is probably a later step after finalizing the conformations of fundamental protein-machines as exemplified by structural conservation of ATP synthase in two distinctly diverge lipid environments of bacteria and archaea^71^.

Our elaboration of lipid-protein chemistry driven amino acid evolution in transmembrane proteins is based on 18:2/18:3 chemistry of mitochondria specific lipid cardiolipin (CL). CL-homeostasis has been implicated in several degenerative conditions including Barth syndrome, cancer, neurodegenerative diseases, non-alcoholic fatty liver disease, and heart failure^72–75^. Unsaturated CL18:3 is so far reported only in plants while 18:2 unsaturation is prevalent in all tissue types of metazoans. However, variation of molar composition of CL 18:2 is reported between tissues within the same organism and even within the same tissue at different metabolic states^32,55,56,76–82^. Our data of instability of human RCs due to further unsaturated CL 18:3 suggest that amino acid composition of trans-IMM helices of RC-subunits are more sensitive to CL-unsaturation than molar composition. However, the diversity of molar composition of 18:2 CL and other lipids may be implicated in the heterogeneous assembly of RCs in different supercomplexes among tissues^55,83,84^ and may partially explain the tissue-specific defects in RC-assembly and function in human diseases with mutations in various RC-subunits^85^. Also, the balance of saturated and unsaturated phospholipids regulates curvature, fluidity, protein stability, and crowding in all cellular membranes^13,86^. Plant lipid metabolism has been evolved to favour polyunsaturated fatty acids (PUFA) which provide membrane plasticity to cope up with diverse habitats^87^.

PUFA is also prevalent in aqueous metazoans to maintain membrane fluidity^88^. Accordingly, investigating evolution of lipid unsaturation in non-mitochondrial membranes and correlated sequence divergence in membrane-embedded proteins would be interesting.

Our results suggest the trans-IMM domain of *Arabidopsis* Ndufa1 is highly unstable and is cleaved after C-terminal asparagine (N) at IMM in human cells. IMM-targeting of proteins is mediated by dedicated import complexes TIM23 and TIM22. These complexes are not water-filled import channels rather their concave surfaces bind and simultaneously expose the clients to IMM-lipids^89–92^. IMM-sorting proteins like Mgr2 or Oxa1 may transiently safeguard the clients from the IMM-lipids after which the clients are either laterally released or conservatively sorted into IMM^93,94^. In both cases, it is imperative that composition of IMM-lipids may directly influence the successful insertion of the clients into IMM. Lipid-assisted insertion mechanism has already been proposed for tail-anchored or signal-anchored proteins at the OMM^95,96^. Our results with Ndufa1 supports lipid-assisted mechanism for IMM-proteins. The mechanism of cleavage of the trans-IMM domain of *Arabidopsis* Ndufa1 in human cells is currently unknown. Membrane-resident serine proteases are known to cleave bonds between N and S^97^. The C-terminal of At-A1 helix also contain consecutive N and S and thus may expose a cleavage sequence for human IMM-proteases after reaching the import channels. Further, there could be impaired binding of the At-A1 helix with the human IMM-sorting proteins Mgr2 or Oxa1 or equivalent import- partners forcing a cleavage. It is also conceivable that the lipid-protein incompatibility may pose significant thermodynamic challenge so that the peptide bond at N is rather cleaved to avoid the energy-expenditure to insert At-A1 helix into human IMM. Addressing these questions will elucidate some thermodynamic puzzles of lipid-protein chemistry.

## Materials and methods

### Analysis of disease-related amino acids in RC-subunits

Online Mendelian Inheritance in Man (OMIM) database (http://omim.org/) was searched for known disease-related mutations in Respiratory Complex (RC) genes. Missense mutations associated with RC-deficiency diseases including Leber hereditary optic neuropathy (LHON) syndrome, Leigh syndrome and mitochondrial encephalomyopathy, lactic acidosis, and stroke-like episodes (MELAS) were considered. Mutations were mapped on available structures of human-RCs (CI – PDB: 5XTD; CII – PDB: 8GS8; CIII – PDB: 5XTE; CIV – PDB: 5Z62; CV – PDB: 8H9V) using PyMol. Bond lengths were measured.

Cut-offs for hydrogen bond – 3.6 Å; salt bridges – 3.9 Å, and cation-π interactions – 5.4 Å. Amino acids with no interactions within these ranges were considered as non-interacting. Surface exposed residues were analysed by commonly used PyMol plug-in FindSurfaceResidues^98–100^ (https://pymolwiki.org/index.php/FindSurfaceResidues). Cut-off for surface exposed residues was increased from default of 2.5 Å^2^ to 3 Å^2^. Red in **Fig. 2D** denotes all the atoms of side-chains being exposed, yellow denotes at least 1 atom of side-chain being exposed, and white denotes completely buried. Conservation analysis of mutating residues was performed using 13 representative organisms from 3 different kingdoms of Eukaryotes i.e., plants, Fungi and Metazoa. Organisms are as follows:

1. Metazoa – *Homo sapiens, Bos taurus, Mus musculus, Danio rerio, Drosophila melanogaster, Caenorhabditis elegans*.
2. Fungi – *Neurospora crassa, Yarrowia lipolytica, Aspergillus nidulans*.
3. Plants – *Arabidopsis thaliana, Zea mays, Hibiscus syriacus, Gracilariopsis chorda*.

Protein sequences of RC-subunits for these organisms were retrieved from NCBI and multiple sequence alignment (MSA) was performed using Clustal Omega from UniPro uGene^101^.

### Structure prediction and alignment

Structures of RC-deficiency disease-mutants of Ndufa2 and Ndufa1 were predicted using RoseTTAFold^102^ (https://robetta.bakerlab.org/). No prediction templates used.

Structures of Ndufa1 chimeras/mutants with or without interacting partners ND1 and Ndufa8 were predicted using AlphaFold2-multimer^103^ (https://colab.research.google.com/github/sokrypton/ColabFold/blob/main/AlphaFold2.ipynb) where parameters were set as default. Predicted structures were aligned to available PDB structure (PDB: 5XTD). RMSDs were calculated using online tool Pairwise Structure Alignment (https://www.rcsb.org/alignment) using jCE alignment method^104^ (9796821) with default parameters.

### blastp search, data filtration and heatmap preparation

Sequences of RC-subunits and subunits of Tom40 complex from organisms such as *Homo sapiens*, *Neurospora crassa*, *Saccharomyces cerevisiae* and *Arabidopsis thaliana* were retrieved from NCBI. blastp^105^ was performed using these sequences where maximum search target was set as 5000 using non-redundant database, BLOSUM62 matrix and rest of the parameters were default. Search results were filtered using in-house python script where sequences corresponding to 134 organisms from Rebeaud et al, 2022^43^ were filtered using e-value cut-off of 0.01 and minimum query coverage of 30% after removing all the duplicate entries. Heatmap was prepared using Morpheus (https://software.broadinstitute.org/morpheus/). *Saccharomyces cerevisiae* sequences were used for blastp searches for Fungi RCs and Tom40 complex. *Neurospora crassa* sequences were used for Fungi CI as *Saccharomyces cerevisiae* lacks canonical CI. Accession number of hits obtained by blastp search are mentioned in **Table S5A-G**.

Violin plot was prepared using blastp filtered results. For a group of subunits, where blast search was restricted to one kingdom, an arbitrary value of 10 was imputed into other kingdoms, to indicate kingdom-specific sequence conservation.

### Mean percent identity calculation of surface-exposed stretches

Continuous stretches of 25 amino acids from representative CI-subunits of *H sapiens* (PDB: 5XTD) and corresponding stretches from *A. thaliana* (PDB: 7ARB) with more than 80% amino acids exposed to surface was identified using FindSurfaceResidues plug-in of PyMol. Amino acid conservation at these exposed stretches was performed by MSA using Clustal Omega from UniPro uGene^101^ on the filtered sequences obtained by blastp across eToL. Maximum number iteration for MSA was set at 100 and rest of the parameters were default. Alignment ProfileGrid measuring % conservation was generated which indicates the % frequency of all 20 amino acids at respective positions^106^. Sequences of surface-exposed stretches of *H. sapiens* and *A. thaliana* were manually compared with the ProfileGrid and % frequencies corresponding to *H. sapeins* and *A. thaliana* sequences were selected. Mean of 25 residues was calculated and was termed as mean % identity. Eight nuclear encoded subunits, each from matrix and IMM arm were used for analysis. Two-sample student t-test was used for significance calculation.

### Simulation

Model 1 and Model 2 IMMs along with trans-IMM helices of Ndufa1 were prepared using CHARMM-GUI server^107–109^. The trans-IMM helices were placed along the bilayer normal (Z-axis) and at the center of the bilayer plane (X-Y plane). Pure IMMs and IMMs with trans-IMM helices were solvated with water molecules and neutralized ions. TIP3P model was used to describe water molecules^110^. KCl was added to neutralize the charge and maintain the physiological salt concentration of 150 mM. Energy minimization was carried out on pure membranes for 5000 steps using the steepest descent method to remove any atomic clashes followed by equilibration with position restraints of lipid headgroups. For model IMMs with trans-IMM helices, energy minimization was carried out on membranes for ∼100 ns, 5000 steps with a short 1ns equilibration with harmonic restrains on trans-IMM helices backbone atoms. Subsequently all restraints were released slowly and the systems were subjected to production run of 1000 ns. First 500 ns was considered as equilibration period and last 500 ns used for trajectory analyses.

Simulations were conducted using the GROMACS tool (version 2021)^111^ and all-atom CHARMM36 force field parameters^112,113^. TIP3P model was used to describe water molecules^110^. The equation of motions was integrated at a timestep of 2 fs, using an isothermal-isobaric (NPT) ensemble. To constrain the covalent bond lengths to hydrogen atoms, a linear constraint solver (LINCS) algorithm^114^ was employed. A Nose−Hoover thermostat was used with a coupling constant of 1.0 ps to maintain the temperature of the systems at 303.15 K^115^. The temperatures of the lipid bilayer and solvent (water and ions) were controlled independently. For models with trans-IMM helices, helix was considered as a separate temperature group. The Parrinello-Rahman barostat was used to control the system’s pressure at 1 bar with a 5.0 ps coupling constant^116^. With a cut-off of 12 Å, the long-range electrostatics interactions are considered via the particle-mesh Ewald (PME) method^117^. The system’s periodic boundary conditions are applied to all three directions within the Ewald summation method. The Lennard-Jones potential was used to compute the van der Waals interactions with a cutoff of 12 Å by employing a force-based switching function (at 10 Å). The analyses were performed using standard gmx tools. VMD was used for trajectory visualization^118^.

5 Å cut-off was used to calculate trans-IMM helix-fatty acyl chain contacts including the only heavy atoms. Number of contacts of all the acyl chains of same lipid species (for example16:0, 18:0, 18:2, 18:3, and 20:4 for PC) were summed together as followed.

Total contacts at matrix leaflet with PC

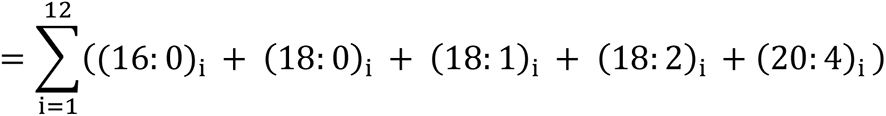

Similarly, number of contacts with other lipids in both leaflets were calculated. Helix-lipid interactions shown in the schematic (**Fig. 3I**) was prepared as per total contacts shown in **Fig. 3H**. Number of lipid molecules per lipid species were placed depending on the total contact that lipid species establishes with the trans-IMM helix. Less than 50 contacts were ignored; 1 lipid molecule was placed for 50-110 contacts; 2 lipid molecules for 110-150 contacts; 3 lipid molecules for 150-160 contacts. Lipid of a particular lipid species that makes maximum contacts with the trans-IMM helix was only use for representation. Rest were ignored.

### Expression constructs

EGFP was PCR-amplified from pCIneo-FlucDM-EGFP^119^ and subcloned in pcDNA4/TO using the restriction enzymes XhoI and XbaI (NEB). Nudfa1 and Ndufa2 was amplified from HEK293T cDNA and cloned into pcDNA4/TO-EGFP using KpnI and XhoI (NEB). Gene sequences of At-A1, Ch-A1 and Hs-A1^Ex-switch^ were codon optimized for human, synthesized, and sub-cloned in pcDNA4/TO-EGFP. Variants of Ndufa2 and Ndufa1 were prepared by site directed mutagenesis using primers in **Methods Table S1**. Ch^Mm-switch^ was prepared by overlap extension PCR, where 1^st^ 23 amino acids were amplified from Neuro2A cDNA and 24^th^ to 70^th^ amino acid was amplified from pcDNA4/TO-Hs-A1-EGFP using primers in **Methods Table S1**. These two fragments were annealed, extended and cloned in pcDNA4/TO-EGFP using Kpn1 and Xho1. 28th to 70^th^ amino acids from Hs-A1 was amplified and inserted before EGFP in pcDNA4/TO-EGFP using Kpn1 and Xho1 to prepare Δ27-Hs-A1-EGFP. Primers are indicated in **Methods Table S1**.

### Cell culture and transfection

HEK293T-6TR (stable overexpression of tetracycline repressor, pcDNA6/TR)^120^ cells were maintained in Dulbecco’s Modified Eagle’s Medium (Gibco) supplemented with 10 % fetal bovine serum (FBS) (Gibco) and 90 U/ml penicillin (Sigma-Aldrich) 50 µg/ml streptomycin (Sigma-Aldrich) at 37°C and 5 % CO2. Transfection of cells was performed with Lipofectamine 3000 reagent (Invitrogen) as per manufacturer’s protocol. For stable cell line preparation, pcDNA4/TO system was used (Life Technologies). HEK293T-6TR cells was transfected with various pcDNA4/TO constructs and selected with 200 µg/ml Zeocin (Invitrogen). Surviving colonies after 2-3 weeks were tested for induction by adding 1 µg/ml Doxycycline (MP Biomedicals) by fluorescence microscopy and western blot. Positive clones were frozen and/or maintained for further experiment.

For western blot and mitochondria isolation, cells were plated and allowed to adhere for 12 hours. 1ug/ml Doxycycline induction was given for 36 hours and cells were pelleted washed with PBS and stored in -80°C until further used. 150mm x 3 dishes was used for mitochondria isolation. 60mm x 1 dish was used for total cell lysate.

### BSA-conjugated lipid supplementation

HEK293T cells stably expressing Hs-A1-EGFP was treated with BSA-conjugated linoleic acid and linolenic acid as described by Omer et al with slight modifications^66^.

Briefly, 5mM solutions of fatty acids (Sigma) were prepared by dissolving in absolute ethanol. Meanwhile, 10% fatty acid free BSA (Sigma) was prepared in 1X PBS (pH 7.4) and incubated at 55°C for 30mins with intermittent vortexing. The solution was cooled to 25°C and filter sterilized. The 5mM fatty acid solution was diluted 1:20 in 10% BSA (final fatty acid concentration: 250μM) and incubated at 55°C for 30mins with intermittent vortexing.

The BSA-conjugated fatty acid stock solution was further diluted 1:10 in incomplete DMEM (final concentration: 25μM of fatty acid in 1% BSA) and supplemented to cells. Control cells were supplemented with BSA-conjugated ethanol. The cells were induced with 1μg/ml doxycycline and incubated at 37°C and 5 % CO2 for 72 hours. Prior lipid supplementation, cells were plated in DMEM and 10% FBS and allowed to adhere for at least 16 hours.

### Immunofluorescence microscopy

Cells were grown on 18mm glass coverslips in 12 well plates (Nunc) and fixed using 4% paraformaldehyde. 0.1% Triton X100 in PBS was used to permeabilize the cells prior to immunostaining. Next, the coverslips were incubated with 5% BSA prepared in PBS with 0.05% Tween-20 (PBST) for 30 min at RT with constant shaking. 1:1000 dilution of Tom20 primary antibody (Santa Cruz Biotechnology) in 0.1% BSA with PBST was added and coverslips were incubated at 4°C overnight followed by PBST wash. Next day, 1:500 dilution of Alexa Fluor Plus 647 Goat anti-Mouse IgG (H+L) Highly Cross-Adsorbed Secondary Antibody (Invitrogen) was added to the coverslips and incubated for 1 hr at room temperature.

Coverslips were mounted on slides using 10 µl Antifade Mounting Medium (cat: H-1000, Vectashield) after DAPI staining. Imaging was done in OLYMPUS FV3000 microscope.

### Mitochondrial fraction preparation, mitoplast preparation, cell lysis, SDS-PAGE, and western blotting

Mitochondria-enriched fraction preparation was performed according to Rawat et al. with slight modification^21^. Briefly, ∼20 million cells were homogenized at 4°C in a Dounce homogenizer with 600 μl of buffer A (83 mM sucrose, 10 mM MOPS pH 7.2). After adding an equal volume of buffer B (250 mM sucrose, 30 mM MOPS pH 7.2), nuclei and unbroken cells were removed by centrifugation at 1000 g for 5 min at 4°C. The supernatant was further centrifuged at 12,000 g for 5 min and washed twice with buffer C (320 mM sucrose, 1 mM EDTA, 10 mM Tris-HCl pH 7.4). The pellet was resuspended in 1X NativePAGE sample buffer (Invitrogen) to obtain mitochondria-enriched fraction followed by storing at -80°C until further use.

Mitoplast preparation protocol was adopted from Hussain M et al. with some modifications^121^. 200 µg of mitochondria-enriched fraction was suspended in hypotonic buffer (20 mM KCl and 10 mM HEPES, pH 7.2) and was treated with trypsin (final concentration 20 µg/ml) (Thermo Scientific) in ice for 15 min. Next, 1X PIC (cocktail of protease inhibitors) (Roche) was added and the fraction was passed through sucrose cushion (0.8 M sucrose in 10 mM HEPES, pH 7.2) at 12000g for 10 min. Pellet fraction was resuspended in hypotonic buffer, 0.1% digitonin, and incubated in ice for 20 min and centrifuged at 12000g. Same steps from resuspension of pellet to passing through sucrose cushion were repeated once with 10 µg/ml trypsin for 5 min to obtain the mitoplasts as pellet.

Cell pellets, mitochondria-enriched pellets, and mitoplast pellets were lysed in SDS-Lysis buffer (0.2 M Tris–HCl pH 6.8, 4 % SDS, 4 % 2-mercaptoethanol, 40 % glycerol), vortexed. Cell pellets were heated at 95°C for 15 min whereas mitochondria-enriched pellets and mitoplast pellets were heated at 55°C for 10 min followed by centrifugation at 12,000 g for 15 min to prepare total cell lysate, mitochondrial fraction, and mitoplast fraction, respectively. Protein estimation was carried out by Amido Black Protein assay, fractions were separated by SDS-PAGE, and transferred onto 0.2 μm PVDF membrane (Roche) for 90 min at 300 mA using Mini-Trans Blot cell system (Bio-Rad). Membranes were probed with primary and secondary antibodies (**Methods Table S2**) and imaged using Chemidoc MP imaging system, Bio-Rad.

### BN–PAGE, *in-gel* activity and native western blotting

Protein content in mitochondria-enriched fractions were estimated using Bradford Assay reagent (Sigma-Aldrich) followed by solubilisation as previously reported by Rawat et al^21^. Briefly, 12 gram digitonin (Sigma-Aldrich) per gram of mitochondria-enriched fraction was added and incubated for 10 min in ice. Supernatant was collected after 30 min centrifugation at 13,000 g. Next, 6 gram of 5% G-250 NativePAGE Sample additive (Invitrogen) was added per gram of mitochondria-enriched fraction. Glycerol was added at final concentration of 10% and the sample was loaded onto NativePAGE 3–12% Bis-Tris Protein Gels (Invitrogen). Native PAGE was run using anode buffer (50 mM Bis-Tris HCl pH 7.0) and blue cathode buffer (50 mM Tricine, 15 mM Bis-Tris HCl pH 7.0, 0.02% Coomassie Brilliant Blue G-250) at 100 V until samples entered the separating gel and then at 150 V until the dye front reached 1/3^rd^ of the entire gel. At this point, blue cathode buffer was replaced with dye-free cathode buffer for the remainder of the run at 250 V until the dye front reached the end of the gel.

*in-gel* CI-Activity – the gel was incubated overnight at room temperature in freshly prepared CI NADH dehydrogenase substrate solution (5 mM Tris-HCl pH 7.4, 0.1 mg/ml NADH and 0.7 mg/ml Nitrotetrazolium Blue chloride). Appearance of purple-bands is indicative of CI activity in the gel. The reaction was stopped with 10% acetic acid, washed with water and imaged.

Native western blotting – Western blotting after BN–PAGE was carried out similar to previously reported^21^. Briefly, the gel was equilibrated in soaking buffer (48 mM Tris-HCl, 39 mM glycine, 0.0037% SDS and 20% methanol) for 30 min and transferred onto 0.45 μm PVDF membrane (Roche) overnight at 30 V, followed by 300 mA for 90 min, using the Mini-Trans Blot cell system (Bio-Rad). The membrane was stained with 0.1% Coomassie Brilliant Blue R-250 dye, 50% methanol and 7% acetic acid and destained with destaining solution 1 (50% methanol and 7% acetic acid) followed by destaining solution 2 (90% methanol and 10% acetic acid) and final destaining using 100% methanol. The membrane was washed with Tris-buffered saline containing Tween 20 (TBST) followed by blocking with 5% BSA in TBST for overnight and probed with appropriate primary and secondary antibodies (**Methods Table S2**).

## Acknowledgement

Dr. Shrish Tiwari, Dr. Mandar V. Deshmukh, Dr. Biswajit Pal, Dr. Hirak Chakraborty for discussion, Debadutta Patra for python script, Pradeep for PyMol analysis. Funding from CSIR grants MLP0143 and MLP0159 is acknowledged.

