## Supplementary figures and methods table for "Kingdom-specific lipid unsaturation shapes up sequence evolution in membrane arm subunits of eukaryotic respiratory complexes"

Figure S1

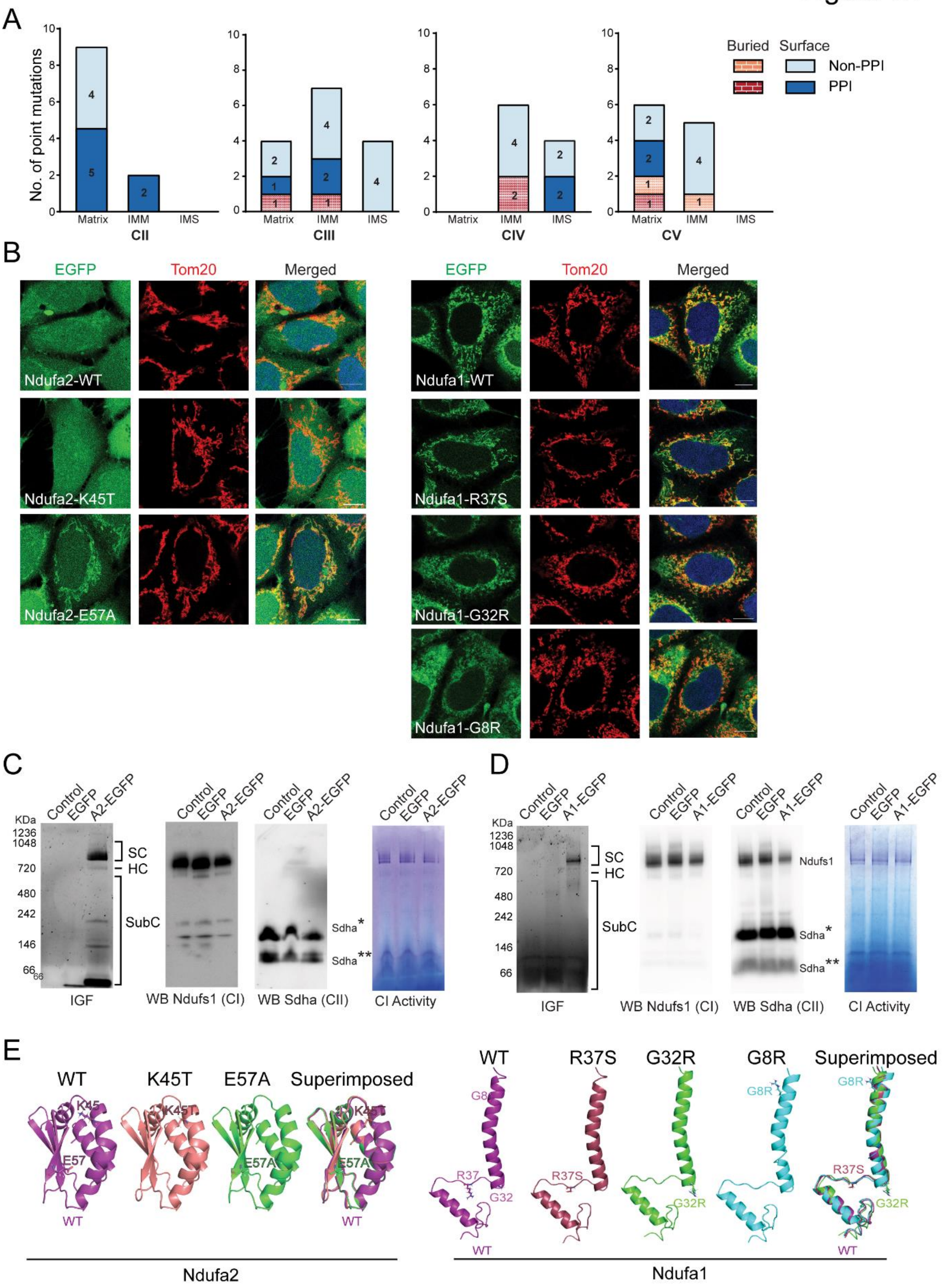

**Fig S1: Mutations causing RC-deficiency diseases are populated at the surface of membrane arm subunits.**

A. Stacked bar graph indicating the distribution of the missense mutations causing RC-deficiency diseases. CII – PDB: 8GS8, CIII – PDB: 5XTE, CIV- PDB: 5Z62, CV-PDB: 8H9V
B. *in-gel* EGFP fluorescence (IGF) assays of overexpressed A2-EGFP, western blots (WBs) with CI and CII subunits using anti-Ndufs1 and anti-Sdha, and CI activity of digitonin solubilised mitochondrial fractions.
C. *in-gel* EGFP fluorescence (IGF) assays of overexpressed A1-EGFP, western blots (WBs) with CI and CII subunits using anti-Ndufs1 and anti-Sdha and CI activity of digitonin solubilised mitochondrial fractions.
D. Different channels of the microscopy images shown in Fig. 1G. Scale bar -10 μm.
E. Homology modelling of WT and mutant Ndufa2 and Ndufa1 using RosseTTAFold. Individual as well as superimposed models are represented. RSMDs between WT and mutants <1.2 Å.

### Figure S2

A

| *H. sapiens*, 5XTD | | | | | *A. thaliana*, 7ARB | | | | *Y.lipolytica*, 7O71 | | | |
|---|---|---|---|---|---|---|---|---|---|---|---|---|
| Subunit | Mutants | Bond length | Interacting subunit | Interacting residue | In solved structure | Bond length | Interacting subunit | Interacting residue | In solved structure | Bond length | Interacting subunit | Interacting residue |
| A12 | R29K | 3.4 | A12 | T65 | Region not resolved | | | | S26 | 2.8 | A12 | K22 |
| S8 | R77W | 2.6 | S2 | E257 | No PPI | | | | R96 | 2.9, 3.2 | S2 | E265 |
| A9 | R360C | 4.4 | ND6 | Y78 | Region not resolved | | | | Region not resolved | | | |
| A6 | R64P | 2.7, 3.1 | AB1 | D111, D114 | S61 | 2.6 | A6 | Y57 | R56 | 2.5, 2.6, 3.6 | AB1 | D65, D68, E71 |
| B3 | W22R | 3 | AB1 | Y85 | Absent | | | | Absent | | | |
| B8 | Y62H | 3 | B8 | D74 | Absent | | | | Absent | | | |

B

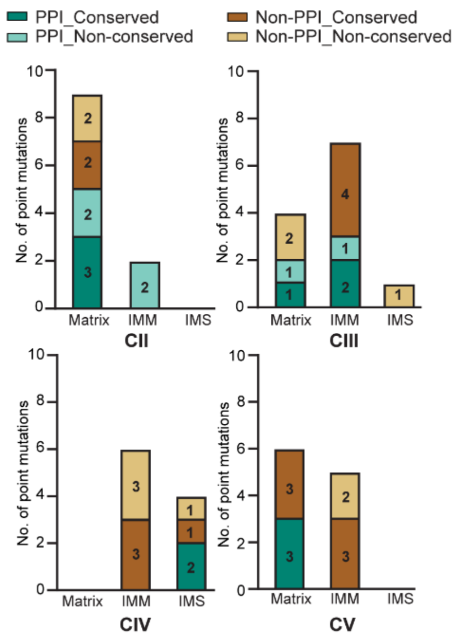


C

| Subunit | S8 | ND1 | ND1 | ND1 | ND1 | ND1 | ND2 | ND2 | B3 | ND5 | ND5 | ND5 | ND5 | ND5 | ND5 |
|---|---|---|---|---|---|---|---|---|---|---|---|---|---|---|---|
| Wild type | E63 | L285 | A52 | E214 | E143 | G131 | L71 | G259 | G70 | A458 | F124 | M237 | D393 | A171 | A236 |
| Physicochemical property | Acidic, Polar | Non-polar | Non-polar | Acidic | Acidic | Non-polar | Non-polar | Non-polar | Non-polar | Non-polar | Aromatic | Non-polar, bulky | Acidic, polar | Non-polar | Non-polar |
| Mutant | Q63 | P285 | T52 | K214 | K143 | S131 | P71 | S259 | T70 | T458 | L124 | L237 | N393 | V171 | T236 |
| Physicochemical property | Zwitter-ionic, polar | Non-polar, Helix breaker | Polar | Basic | Basic | Polar | Non-polar, Helix breaker | Polar | Polar | Polar | Aliphatic | Non-polar, less bulky | Zwitter-ionic, polar | Non-polar, bulky | Polar |

D

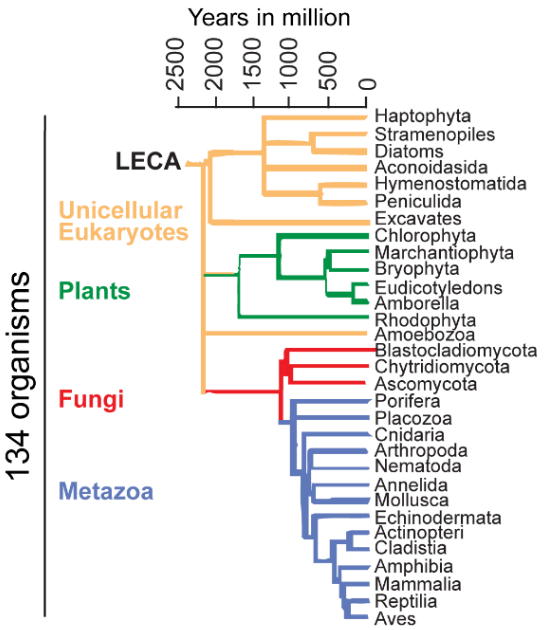


Adapted from Rebeaud et al, 2021

E

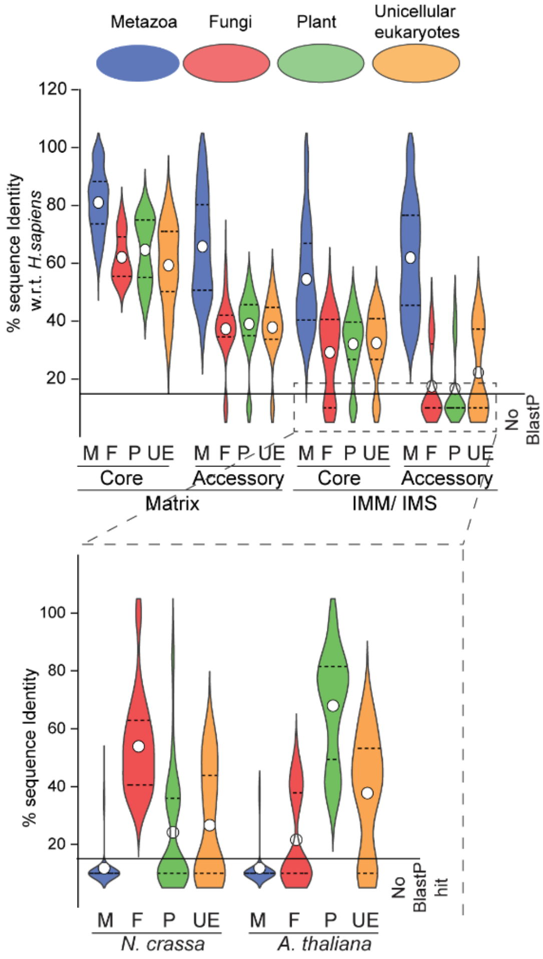

**Fig S2: RC-deficiency disease mutations in IMM-arms are less conserved across eukaryotes**

A. Table indicating unconserved matrix exposed amino acids involved in PPI in *H. sapiens* (PDB: 5XTD) and associated with CI-deficiency diseases are shown. Corresponding amino acids in *A. thaliana* (PDB: 7ARB) and *Y. lipolytica* (PDB: 7O71) with their interacting residues indicated. Bond length: Å. Region not resolved: the amino acid is present but not resolved in published Cryo-EM structure; Stretch not present: absent in that organism.

B. Stacked bar graph representing conservation of missense mutations involved in CII, CIII, CIV and CV deficiency diseases. Thirteen organisms from 3 different kingdoms as shown in **2A** were considered for analysis.

C. Table indicating conserved exposed residues of CI IMM arm with changes in physicochemical properties upon disease mutation. Amino acids are indicated by 1 letter code followed by position of the amino acid

D. Eukaryotic tree of life (ToL) adapted from Rebeaud et al, 2021[1]. 134 organisms from different kingdoms shown that are considered for blastp data analysis shown in **2E**.

E. Violin plot indicating the mean % identity and the spread of mean % identity analysis shown in **2E**. **Top:** mean % identity of all CI-subunits against *H. sapiens* sequence. **Bottom:** mean % identity analysed against *N. crassa*, *A. thaliana* sequences for IMM arm subunits with no blastp hits indicated above.  Circle – mean; dashed lines – quartiles.

Figure S3

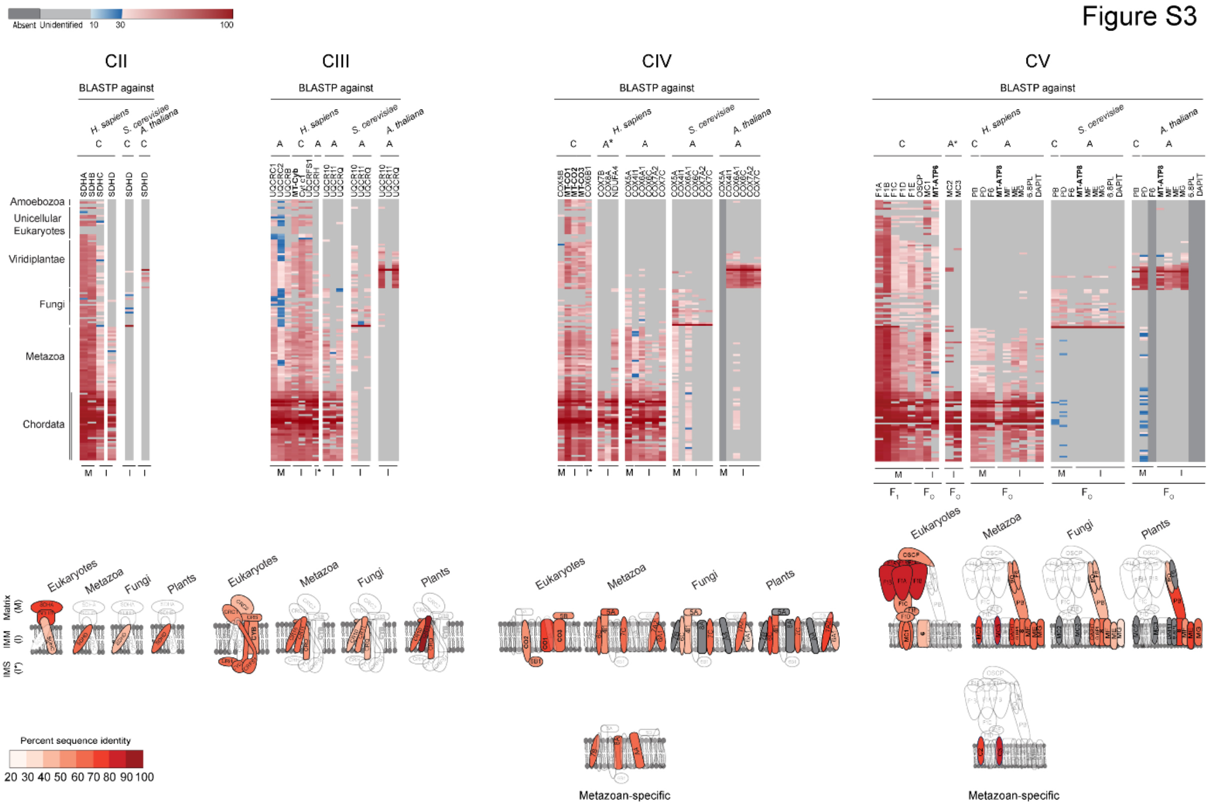


M - MATRIX, I - IMM, I* - IMS, C - Core, Core, A - Accessory, A* - Metazoan-specific Accessory
**Mito encoded**

**Fig S3: IMM-arm subunits of RCs are sequence conserved within eukaryotic kingdom**

A. **Top:** Heatmaps representing % sequence identity of individual subunits of CII, CIII, CIV and CV when searched using blastp. Sequences of *Homo sapiens, Saccharomyces cerevisiae and Arabidopsis thaliana* from Metazoa, Fungi and plants were used for blast search. Light grey indicates no blastp hit found. Dark grey indicates subunit absent in particular kingdom. M- matrix; I- IMM; I*- IMS; C - Core; A – Accessory. For CV, subunits are prefixed as ATP5 except for MT-ATP6 and MT-ATP8. **Bottom:** Schematic representations indicating the mean % identity of individual subunits.

Figure S4

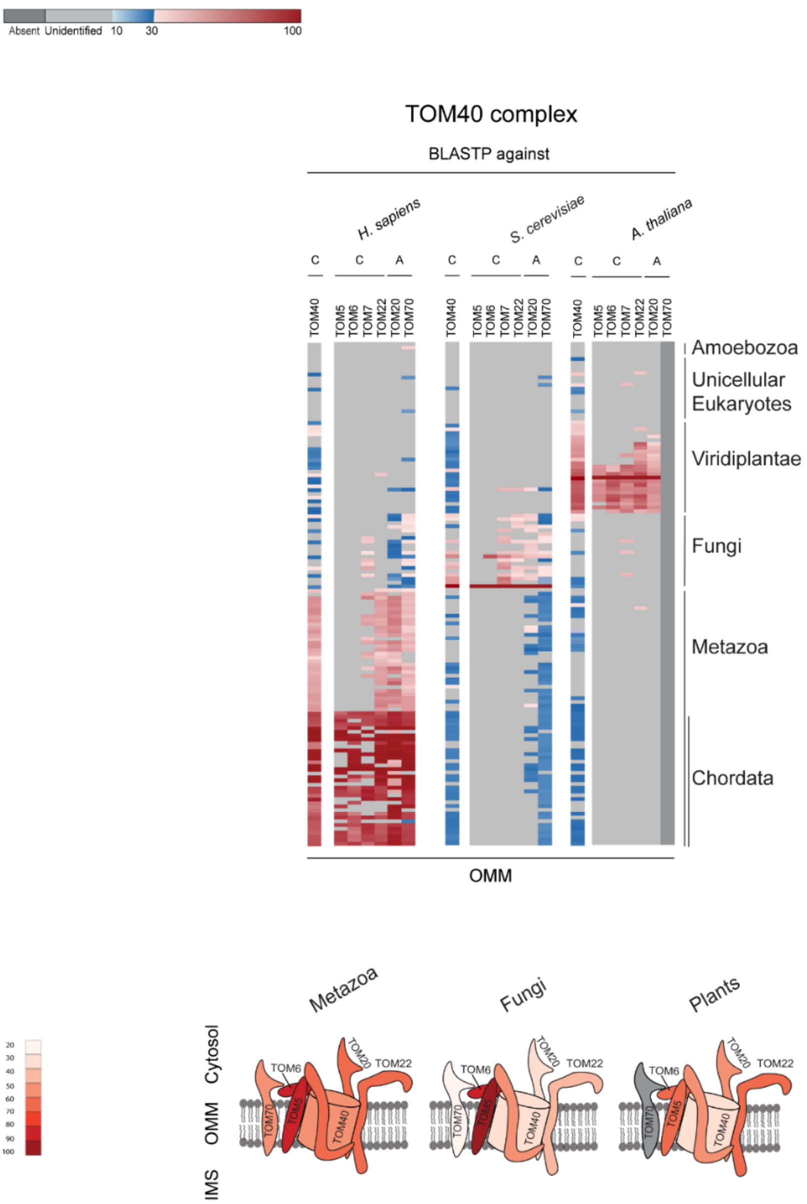


**Fig S4: Membrane-arm subunits of Tom40 complex are sequence conserved within eukaryotic kingdom**

A. **Top:** Heatmaps representing % sequence identity of individual subunits of OMM complex Tom40 when searched using blastp. Sequences of *Homo sapiens, Saccharomyces cerevisiae and Arabidopsis thaliana* from Metazoa, Fungi and plants were used for blast search. Light grey - no blastp hit found. Dark grey - subunit is absent in particular kingdom. C- Core; A- Accessory. **Bottom:** Schematic representation indicating the mean % identity of individual subunits.

Figure S5

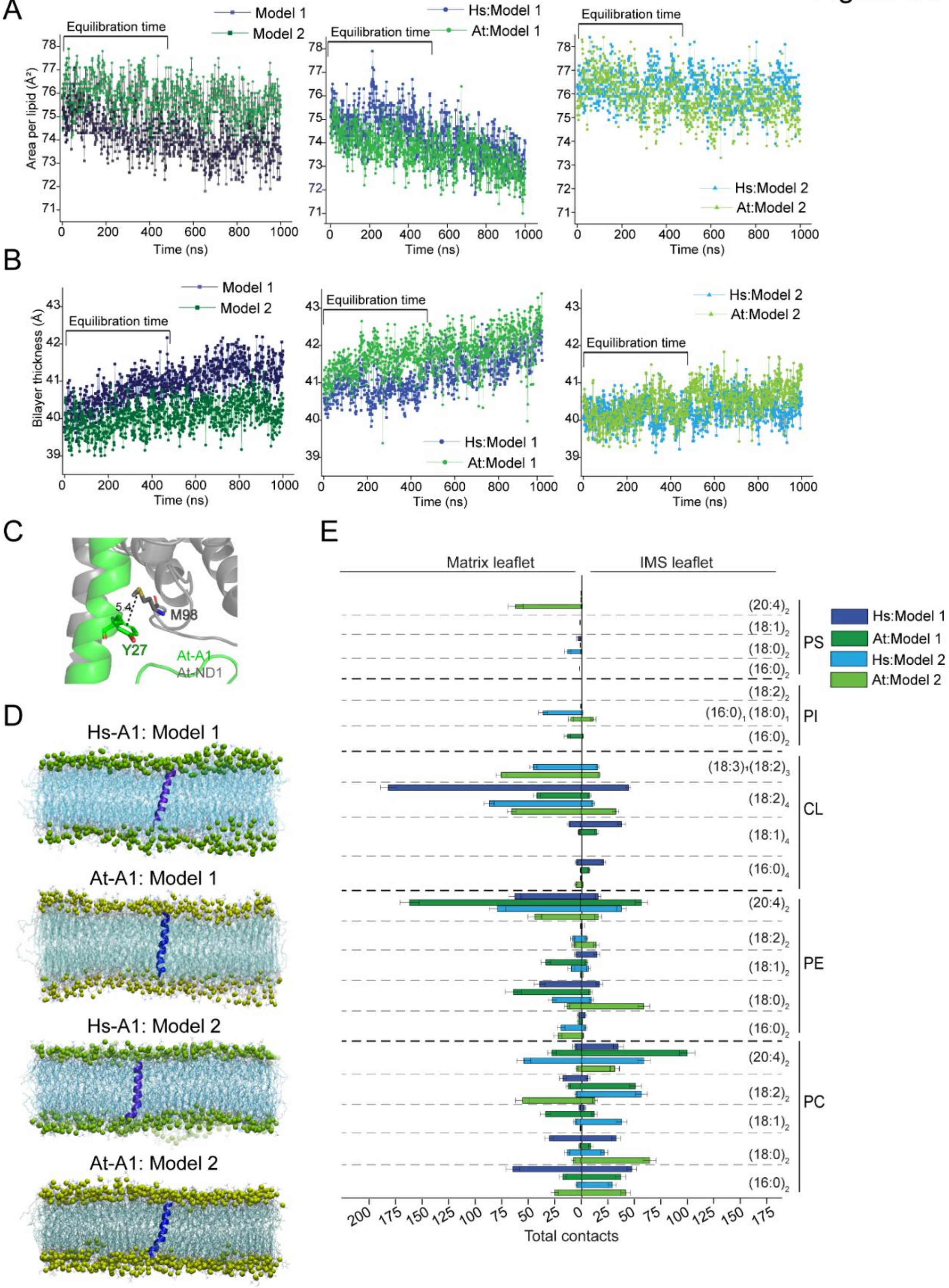

**Fig S5: Trans-IMM helix of Human and Arabidopsis Ndufa1 interacts differently with IMM-lipids**

A. Line graph indicating area per lipid ($Å^2$) across simulation time (1 μs). **Left:** Model membrane 1 and 2 without trans-IMM helix. **Middle:** At-A1 (At) and Hs-A1 (Hs) trans-IMM helix in model 1. **Right:** At-A1 (At) and Hs-A1 (Hs) trans-IMM helix in model 2.
B. Line graph indicating bilayer thickness (Å) across simulation time (1 μs). **Left:** Model membrane 1 and 2 without trans-IMM helix. **Middle:** At-A1 (At) and Hs-A1 (Hs) trans-IMM helix in model 1. **Right:** At-A1 (At) and Hs-A1 (Hs) trans-IMM helix in model 2.
C. Stick representation of Y27 of Ndufa1 forming cation-π interaction with M98 of ND1 in *A. thaliana* as visualized by PyMol. Dashed black lines indicating interaction with bond length in Å. PDB: 7ARB.
D. Simulation snapshots of Hs-A1 (purple) trans-IMM helix in Model 1 and Model 2 (Phosphate plane: green) and At-A1 (blue) trans-IMM helix in in Model 1 and Model 2 (Phosphate plane: lime green). Acyl chains – light blue.
E. Graph showing time-average total acyl chains of contacts of different lipid species of matrix and IMS leaflets with Ndufa1 trans-IMM helices i.e., human M1-M12, *A. thaliana* V4-I15 and human G13-T23, *A. thaliana* G16-Q26, respectively. Total contacts averaged over final 500 ns of simulation time. Error bar: SD.

Figure S6

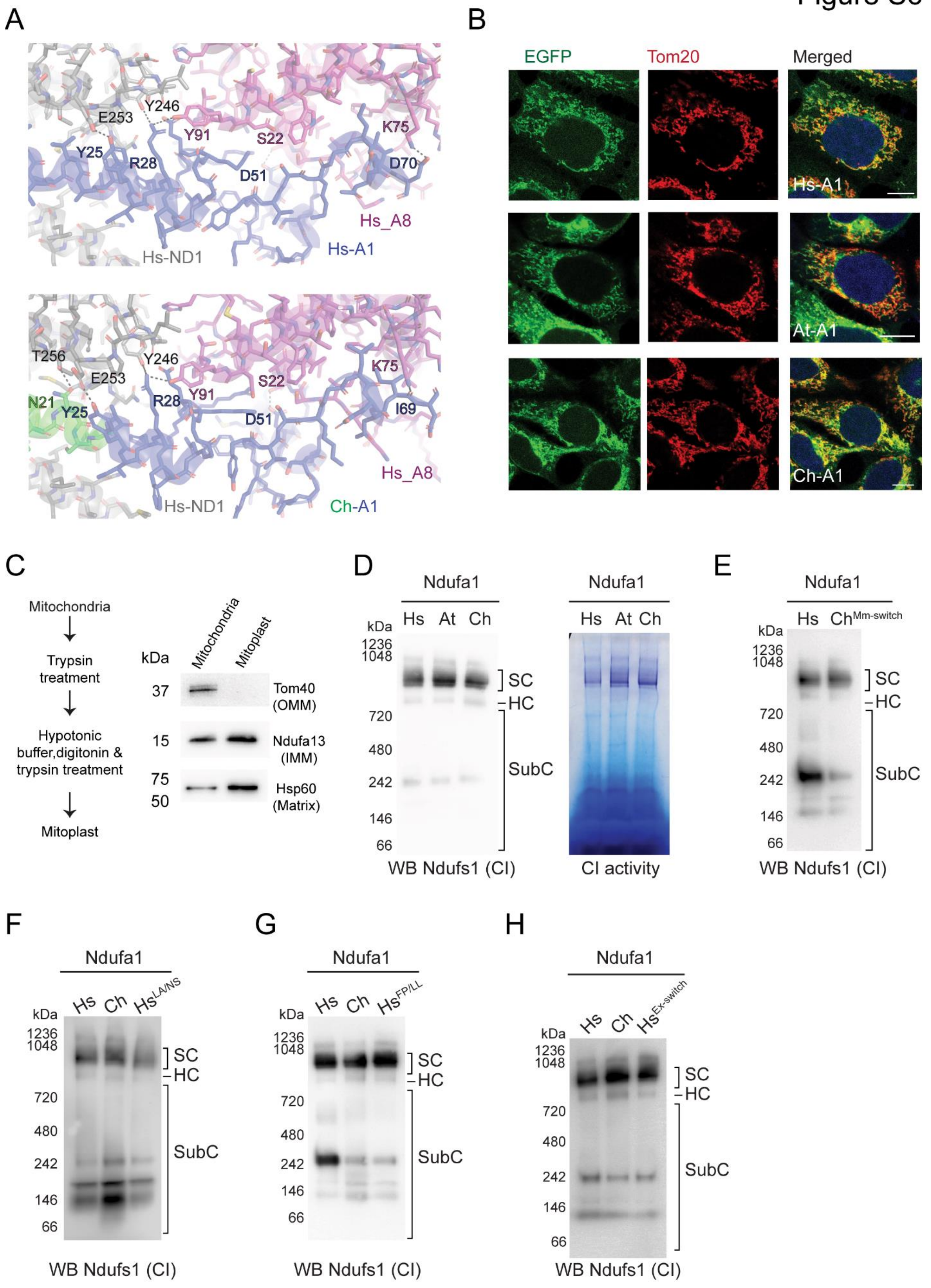

A
E253
Y246
Y91
S22
K75
Y25
R28
D51
D70
Hs_A8
Hs-ND1
Hs-A1
T256
E253
Y246
K75
N21
Y25
R28
Y91
S22
D51
I69
Hs_A8
Hs-ND1
Ch-A1
B
EGFP
Tom20
Merged
Hs-A1
At-A1
Ch-A1
C
Mitochondria
Trypsin treatment
Hypotonic buffer,digitonin & trypsin treatment
Mitoplast
kDa
Mitochondria
Mitoplast
37
Tom40 (OMM)
15
Ndufa13 (IMM)
75
50
Hsp60 (Matrix)
D
Ndufa1
Hs At Ch
kDa
1236
1048
720
480
242
146
66
SC
HC
SubC
WB Ndufs1 (CI)
Ndufa1
Hs At Ch
CI activity
E
Ndufa1
Hs Ch^Mm-switch
kDa
1236
1048
720
480
242
146
66
SC
HC
SubC
WB Ndufs1 (CI)
F
Ndufa1
Hs Ch Hs^LA/NS
kDa
1236
1048
720
480
242
146
66
SC
HC
SubC
WB Ndufs1 (CI)
G
Ndufa1
Hs Ch Hs^FP/LL
kDa
1236
1048
720
480
242
146
66
SC
HC
SubC
WB Ndufs1 (CI)
H
Ndufa1
Hs Ch Hs^Ex-switch
kDa
1236
1048
720
480
242
146
66
SC
HC
SubC
WB Ndufs1 (CI)

**Fig S6: Switching lipid-exposed amino acids of Hs-A1 trans-IMM helix with At-A1 trans-IMM helix renders *in-cellulo* incompatibility**

A. **Top:** Stick representation indicating H-bonds observed between Hs-A1, Ndufa8 and ND1 (PDB:5XTD). **Bottom:** H-bonds observed between Ch-Ndufa1, Hs-A8 and Hs-ND1 as visualized by PyMol. Hs-ND1:grey; Hs-A8:pink; Hs-A1:blue; Ch-A1:blue and green; Dashed black lines indicating interaction.
B. Microscopy images showing EGFP tagged Hs-A1, At-A1, and Ch-A1. Each image represent single Z-section of 0.3 μm. DAPI (blue): nucleus. Tom20: mitochondrial marker. Scale bar: 10 μm.
C. **Left:** Schematic explaining preparation of mitoplast. **Right:** Immunoblot of mitochondrial fraction and mitoplast fraction. Tom40 - Outer mitochondrial membrane marker; Ndufa13 - Inner mitochondrial membrane marker; Hsp60 - matrix marker
D. **Left:** Western blot (WB) for CI subunit Ndufs1 using digitonin solubilised mitochondrial fractions run on native PAGE from cells shown in **4H**. SC- supercomplex; HC- holocomplex; SubC- subcomplex. **Right:** *in-gel* CI activity.
E. Western blot (WB) for CI subunit Ndufs1 using digitonin solubilised mitochondrial fractions run on native PAGE from cells shown in **4I**. SC- supercomplex; HC- holocomplex; SubC- subcomplex.
F. Western blot (WB) for CI subunit Ndufs1 using digitonin solubilised mitochondrial fractions run on native PAGE from cells shown in **5D**. SC- supercomplex; HC- holocomplex; SubC- subcomplex.
G. Western blot (WB) for CI subunit Ndufs1 using digitonin solubilised mitochondrial fractions run on native PAGE from cells shown in **5E**. SC- supercomplex; HC- holocomplex; SubC- subcomplex.
H. Western blot (WB) for CI subunit Ndufs1 using digitonin solubilised mitochondrial fractions run on native PAGE from cells shown in **5G**. SC- supercomplex; HC- holocomplex; SubC- subcomplex.

**Methods table S1: Primers used for cloning**

| Gene name | Primer | Primer sequence | Source |
|---|---|---|---|
| EGFP | Forward | CTCGAGATGGTGAGCAAGGGCGAGGAG | Bioserve |
| | Reverse | CCTCTAGATTACTTGTACAGCTCGTCCATG | Bioserve |
| Ndufa2 | Forward | GGTACCATGGCGGCGGCCGCAGCAA | Bioserve |
| | Reverse | CTCGAGGGCTTTACCACTTAGAACGTTCTCCAGGGC | Bioserve |
| Ndufa2-K45T | Forward | GCTACGTGGAGCTGACAAAGGCGAATCCCGACCTAC | Bioserve |
| | Reverse | GTAGGTCGGGATTCGCCTTTGTCAGCTCCACGTAGC | Bioserve |
| Ndufa2-E57A | Forward | CCATCCTAATCCGCGCTTGCTCCGATGTGCAGC | Bioserve |
| | Reverse | GCTGCACATCGGAGCAAGCGCGGATTAGGATGG | Bioserve |
| Ndufa1 | Forward | GGTACCATGTGGTTCGAGATTCTCCC | Bioserve |
| | Reverse | CTCGAGATCAATGTTCTCCAAACCCTTTG | Bioserve |
| Ndufa1-G32R | Forward | CACAGGTTCACTAACAGGGGCAAGGAAAAAAGGG | Bioserve |
| | Reverse | CCCTTTTTTCCTTGCCCCTGTTAGTGAACCTGTG | Bioserve |
| Ndufa1-R37S | Forward | GGGGCAAGGAAAAAAGCGTTGCTCATTTTGGG | Bioserve |
| | Reverse | CCCAAAATGAGCAACGCTTTTTTCCTTGCCCC | Bioserve |
| Ndufa1-G8R | Forward | GGTTCGAGATTCTCCCCAGGCTCTCCGTCATGGG | Bioserve |
| | Reverse | CCCATGACGGAGAGCCTGGGGAGAATCTCGAACC | Bioserve |
| Hs-Ndufa1$^{LA/NS}$ | Forward | GCTTGTTGATTCCAGGAAACAGCACTGCGTACATCCAC | Bioserve |
| | Reverse | GTGGATGTACGCAGTGCTGTTTCCTGGAATCAACAAGC | Bioserve |
| Hs-Ndufa1$^{FP/LL}$ | Forward | GTGGCTGGAGATTCTCCTGGGACTCTCCGTCATGGG | Bioserve |
| | Reverse | CCCATGACGGAGAGTCCCAGGAGAATCTCCAGCCAC | Bioserve |
| Ch-Ndufa1-N21L | Forward | CTGTGCATCATGGGCCTGAGCCAGGCGTACATCC | Bioserve |
| | Reverse | GGATGTACGCCTGGCTCAGGCCCATGATGCACAG | Bioserve |
| Ch-Ndufa1-Q23T | Forward | GGCATGCTGTGCATCATGGGCAACAGCACTGCGTACAT CCACAGG | Bioserve |
| | Reverse | GTGAACCTGTGGATGTACGCAGTGCTGTTGCCCATGAT GCACAGC | Bioserve |
| Ch$^{mm\text{-}switch}$ | Forward | GGTACCATGGCGGCGGCCGCAGCAA | Bioserve |
| Ch$^{mm\text{-}switch}$-Mm-overlap | Reverse | TGTGGATGTACGCAGTGGACACCC | Bioserve |
| Ch$^{mm\text{-}switch}$-Hs-overlap | Forward | ACTGCGTACATCCACAGGTTCACTAACGG | Bioserve |
| Ch$^{mm\text{-}switch}$ | Reverse | CTCGAGGGCTTTACCACTTAGAACGTTCTCCAGGGC | Bioserve |
| Δ27-Hs-Ndufa1 | Forward | GGTACCATGAGGTTCACTAACGGGGGCAAGGAAA | Bioserve |
| | Reverse | CTCGAGATCAATGTTCTCCAAACCCTTTG | Bioserve |

#### Methods table S2: Antibodies used for western blots and ICC

| Antibody name | Company | Catalog number | Dilution | |
|---|---|---|---|---|
| | | | **ICC** | **Western** |
| **Primary Antibody** | | | | |
| Chicken polyclonal anti-GFP Antibody | Abcam | ab13970 | N/A | 1:5000 |
| Rabbit monoclonal Anti-Ndufs1 antibody [EPR11521(B)] | Abcam | ab169540 | N/A | 1:1000 |
| Mouse monoclonal anti SDHA Antibody (2E3GC12FB2AE2) | Abcam | ab14715 | N/A | 1:2000 |
| Mouse monoclonal Anti-Hsp60 antibody [LK-1] | Abcam | ab59457 | N/A | 1:5000 |
| Mouse monoclonal, Beta Actin antibody | Abcam | ab8224 | N/A | 1:10000 |
| Mouse monoclonal Anti-Tom20 Antibody (F10) | Santa Cruz Biotechnology | sc-17764 | 1:1000 | N/A |
| Mouse monoclonal Anti-Grim19 antibody [6E1BH7] (Ndufa13) | Abcam | ab110240 | N/A | 1:500 |
| Mouse monoclonal Anti-Tom40 antibody | Santa Cruz Biotechnology | sc-365467 | N/A | 1:500 |
| Rabbit monoclonal Anti-Glud1+Glud2 antibody [EPR11369B] | Abcam | ab166618 | N/A | 1:2000 |
| **Secondary Antibody** | | | | |
| Anti-Rabbit IgG (whole molecule) (peroxidase conjugate) | Sigma | A6154 | N/A | 1:20000 |
| Anti-Mouse IgG (whole molecule) (peroxidase conjugate) | Sigma | A4416 | N/A | 1:20000 |
| Goat anti-Mouse IgG (H+L) Highly Cross-Adsorbed Secondary Antibody, Alexa Fluor Plus 647 | Invitrogen | A32728 | 1:500 | N/A |
| Donkey anti-Chicken IgY (H+L) Cross-Adsorbed Secondary Antibody, HRP | Invitrogen | SA1-300 | N/A | 1:7500 |
